# POMBOX: a fission yeast toolkit for molecular and synthetic biology

**DOI:** 10.1101/2023.05.24.542151

**Authors:** Téo Hebra, Helena Smrčková, Büsra Elkatmis, Martin Převorovský, Tomáš Pluskal

**Affiliations:** Institute of Organic Chemistry and Biochemistry of the Czech Academy of Sciences, Praha, Czech Republic; Department of Cell Biology, Faculty of Science, Charles University, Prague, Czech Republic

## Abstract

*Schizosaccharomyces pombe* is a popular model organism in molecular biology and cell physiology. With its ease of genetic manipulation and growth, supported by in-depth functional annotation in the PomBase database and genome-wide metabolic models, *S. pombe* is an attractive option for synthetic biology applications. However, *S. pombe* currently lacks modular tools for generating genetic circuits with more than one transcriptional unit. We have developed a toolkit to address this issue. Adapted from the MoClo-YTK plasmid kit for *Saccharomyces cerevisiae* and using the same Golden Gate grammar, our POMBOX toolkit is designed to facilitate the fast, efficient and modular construction of genetic circuits in *S. pombe*. It allows for interoperability when working with DNA sequences that are functional in both *S. cerevisiae* and *S. pombe* (e.g. protein tag, antibiotic resistance cassette, coding sequences). Moreover, POMBOX enables the modular assembly of multi-gene pathways and increases possible pathway length from 6 to 12 transcriptional units. We also adapted the stable integration vector homology arms to Golden Gate assembly and tested the genomic integration success rate depending on different sequence sizes, from four to twenty-four kilobases. We included fourteen *S. pombe* promoters that we characterized for two fluorescent proteins, in both minimal defined media (EMM2) and complex media (YES). Then we tested six *S. cerevisiae* and six synthetic terminators in *S. pombe*. Finally, we used the POMBOX kit for a synthetic biology application in metabolic engineering and expressed plant enzymes in *S. pombe* to produce specialized metabolite precursors, namely methylxanthine, amorpha-4,11-diene and cinnamic acid from the purine, mevalonate and amino acid pathways.

## Introduction

The fission yeast *Schizosaccharomyces pombe* is a well-characterized model organism for molecular and cellular biology of eukaryotes.^1^ As a yeast, *S. pombe* is a unicellular archiascomycete that grows to high density and is easy to manipulate. Its extensive experimental functional annotations have been compiled in the well-curated database PomBase.^1, 2^ Those assets make *S. pombe* an attractive organism for synthetic biology where the ideal chassis organism has to be easy to genetically modify and its behavior precisely predicted and modeled. Among the applications of synthetic biology, metabolic engineering aims to increase the production of high-value chemicals through the modification of organisms that naturally produce these compounds^3^ or by expressing a pathway of interest in a heterologous host^4–6^

Evolutionarily, *S. pombe* diverged from *S. cerevisiae* and other yeasts (*Candida*, *Yarrowia*, *Pichia* spp.) about one billion years ago,^7^ and has several characteristics that make it a potential chassis for metabolic engineering. Basic molecular biology tools, such as reporter genes, the CRISPR/Cas system, and genomic integration, are already available for use with *S. pombe*. Additionally, genome-scale metabolic models have been established for this organism. Unlike *S. cerevisiae*, *S. pombe* retains 4′-phosphopantetheinyl transferase, which is necessary for the synthesis of fungal and bacterial polyketides and non-ribosomal peptides,^8^ and produces cofactors required for specific enzymatic reactions, such as vitamin B21. Some projects have proposed using *S. pombe* as a metabolic engineering platform, particularly for overproducing 3-hydroxypropionic acid via the malonyl-CoA pathway,^9^ lactic acid,^10^ ricinoleic acid^11^ and vanillin,^12^ or for expressing cytochrome P450 with its partner NADPH-cytochrome P450 oxidoreductases (CPRs).^13^ However, these few examples are mostly limited to the expression of a single heterologous enzyme, or at most, in the case of vanillin, a biosynthetic pathway consisting of three enzymes. To recreate the vanillin biosynthetic pathway, Hansen *et al.* had to sequentially integrate the three genes into three different loci. The absence of tools for performing metabolic engineering in *S. pombe* has been identified as a factor contributing to the lack of non-ribosomal peptide production utilizing this organism.^8^

Synthetic biology has greatly contributed to metabolic engineering through the improvement of genome editing^14^ and multi-gene^15^ assembly tools for the efficient reconstruction and integration of biosynthetic pathways. Promoter and terminator libraries^16, 17^ provide finely characterized regulatory elements and minimize construct size and homologous recombination events to ensure the stability and tunable expression of biosynthetic pathways. However, none such tools currently exists for *S. pombe*. For plasmid or integration vector construction, efforts have been made to develop strategies that allow for the modularity of promoter and coding sequences. A toolkit based on Golden Gate assembly, a fast and efficient DNA assembly method to ligate multiple DNA fragments in a single reaction,^18^ was proposed by Kakui *et al*. in 2015^19^ but only provided three different promoters (*adh1*, *nmt1* and *urg1*) and a single terminator. Furthermore, this system was not designed for the construction of multiple gene circuits. In 2020, Vještica *et al*. proposed a series of new integration vectors for *S. pombe* making the regulatory elements modular.^20^ The Stable Integration Vectors (SIVs) they built allow for efficient and rapid integration of DNA sequences into *S. pombe* genome. They also characterized six new promoters and used the exogenous terminator tCyc1 from *S. cerevisiae*. Therefore, there is currently no solution for the fast and modular multi-gene assembly and reconstruction of biosynthetic pathways in *S. pombe*. In 2015, Lee *at al*.^21^ designed the molecular cloning yeast toolkit (MoClo-YTK) for *S. cerevisiae*. It contains 96 characterized parts split into eight part types (i.e. connectors left and right, promoter, coding sequence, terminator, yeast marker, origin of replication, bacterial marker) enabling the streamlined assembly of cassette and multi-gene plasmids in a modular fashion. Thanks to GoldeGate assembly and its standardized overhangs, this toolkit has been extended for other applications and adapted to other organisms.^22–28^

Here, we introduce POMBOX, a toolkit dedicated to the modular assembly of multi-gene integration vectors for applications in molecular and synthetic biology. POMBOX reuses the overhangs proposed in the MoClo-YTK^21^ toolkit and is therefore compatible with several other existing molecular biology toolkits, allowing for better interoperability between DNA parts. We also include short synthetic regulatory elements to decrease the size of constructs and maximize the stability of the constructs after genomic integration.

## Results

### Principles of the toolkit

POMBOX is an extension of MoClo-YTK designed for *S. pombe*. The principle of the toolkit is briefly described here. An extensive explanation, details of the workflow and examples are presented in the supplementary information and in the original MoClo-YTK article.^21^ Golden Gate assembly is in principle based on type IIS restriction enzymes. Unlike classical restriction enzymes, type IIS restriction enzymes cleave DNA outside the recognition site (Figure 1A). The Golden Gate approach offers two significant advantages for molecular biology strategies. First, positioning the recognition sites outside of DNA sequences to be cloned makes it possible to generate products lacking the original restriction site (Figure 1A). Second, as the overhangs can consist of any sequence of four nucleotides, they can be designated upstream to enhance ligation efficiency,^29, 30^ resulting in a Golden Gate "grammar" that enables reliable ligations (Figure 1B). To harness these properties, the MoClo-YTK toolkit adopts the Golden Gate assembly approach, defining eight types of DNA parts essential for generating plasmids or genomic integration vectors. Assembly connectors (Parts 1 and 5) facilitate genotyping and multi-gene assembly. Parts 2, 3 and 4 form the transcription unit, constituted by a promoter (Part 2), a coding sequence (Part 3) and a terminator (Part 4). Additionally, subtypes of Parts 3A, 3B, 4A and 4B have been developed to integrate tags to the coding sequence at the N-terminus or C-terminus (Figure S1). These various parts are then employed in a streamlined one-step Golden Gate reaction (Figure 1D), enabling the assembly of a complete functional plasmid suitable for experimentation or the generation of a multi-gene plasmid (Figure 1E). To function properly, some core rules have to be followed when generating new parts: (I) DNA sequences should be free of BsaI, BsmBI and NotI recognition sites, as those three enzymes are used for plasmid assembly and vector linearization; and (II) Overhangs should respect the MoClo-YTK grammar. Overhangs are listed in Supplementary Information, Tutorial section.

**Figure 1:**
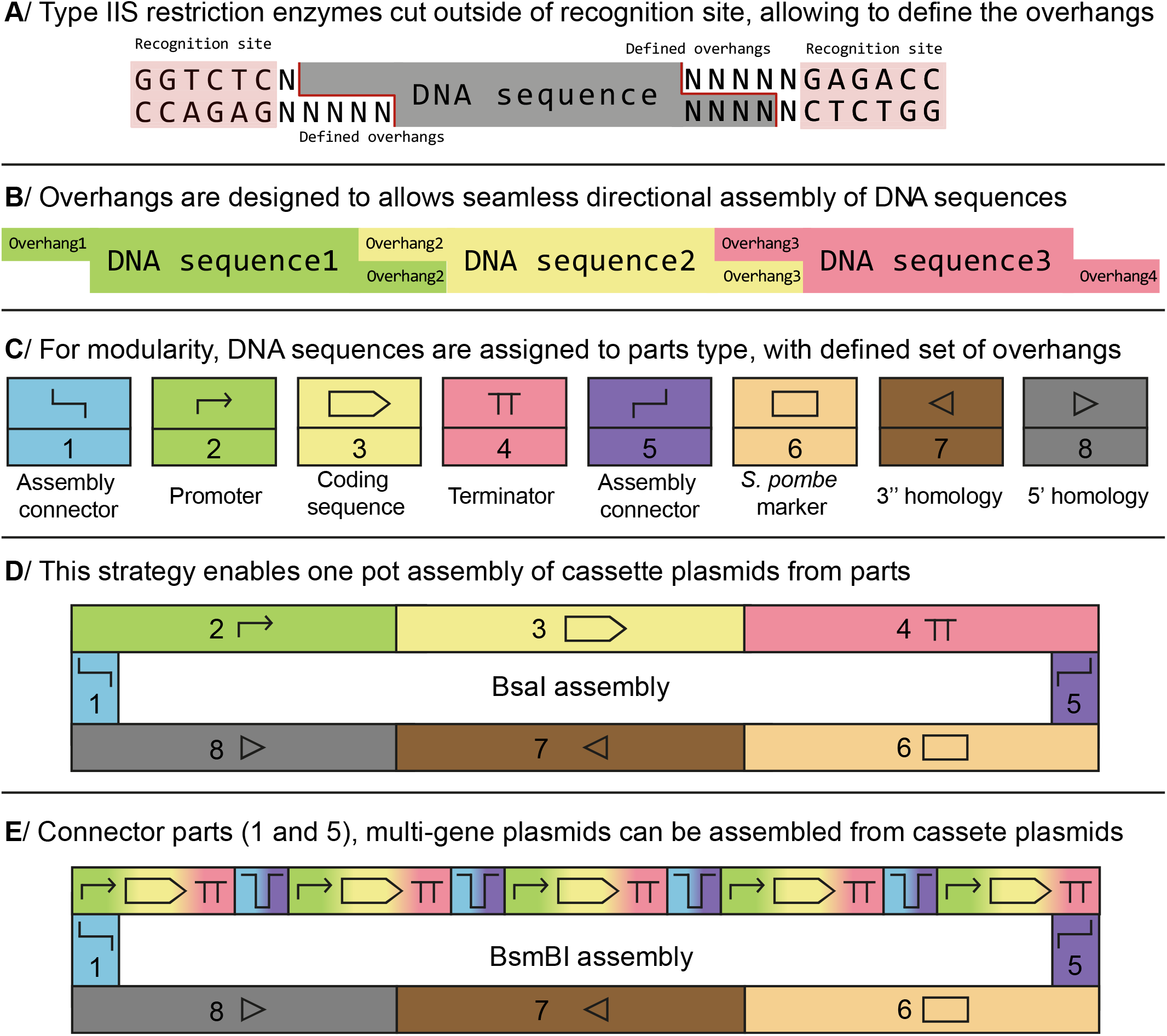
Standardized Golden Gate assembly workflow for plasmid assembly. (**A**) Golden Gate assembly relies on type IIS restriction enzymes. When the recognition sites are outside the DNA sequence of interest, it allows for seamless assembly. (**B**) Overhangs can be defined to increase ligation efficiency and assemble DNA in an ordered fashion. (**C**) The toolkit relies on eight part types, each one with a defined set of overhangs. It enables the integration of new compatible parts in the toolkit. (**D**) Parts are used to assemble, in one pot, a complete plasmid or integration vector, using the BsaI enzyme. It enables modularity of sequences and combinatorial assembly. (**E**) Pairs of connector parts allow for the assembly of multi-gene plasmids using the BsmBI enzyme.

Overall, this molecular biology framework enables the fast, efficient and reliable generation of plasmids. All the backbone vectors used in this toolkit possess a fluorescent protein drop-out allowing for fast selection of transformants with a correctly assembled plasmid thanks to green-white screening. As claimed in the MoClo-YTK paper,^21^ screening one transformant is sufficient to find a correctly assembled transformant plasmid in most cases. Using POMBOX and the MoClo-YTK assembly principle, we were able to generate up to 24 strains of *S. pombe* expressing a fluorescent protein under the control of different transcriptional regulators in seven days.

POMBOX is a collection of characterized DNA parts (**Figure 2A**), based on the Golden Gate assembly grammar of Lee *et al*. We chose this Golden Gate grammar because some exogenous sequences (antibiotic resistance cassettes, bacterial markers and origins of replication, coding sequences, tags, and assembly connectors; see **Figure S1** for a list of compatible parts from the MoClo-YTK toolkit) can be shared between *S. cerevisiae* and *S. pombe*. To complete the toolkit, we propose forty new parts and two integration vectors (pPOM001-042): six new pairs of connectors, fourteen promoters characterized in two different *S. pombe* culture media and for two protein expressions. Then we verified the compatibility of *S. cerevisiae* terminators and short synthetic terminators in *S. pombe*, and adapted the Vještica *et al*.^20^ strategy for stable genomic integration in *S. pombe* for compatibility with the Golden Gate toolkit. We assessed the impact of the exogenous DNA length of the transformation rate of *S. pombe*. POMBOX DNA parts can be used and adapted for regular molecular biology application such as protein expression (**Figure 1B**, example 1), epitope tagging (**Figure 2B**, example 2), gene deletion, insertion of mutations, and protein–protein interactions. The procedure to generate the two examples of integration vectors from POMBOX is described in the supplementary data. Finally, to validate the relevance of POMBOX, we chose metabolic engineering applications and expressed three plant enzymes building precursors for the biosynthesis of specialized metabolites in *S. pombe*.

**Figure 2:**
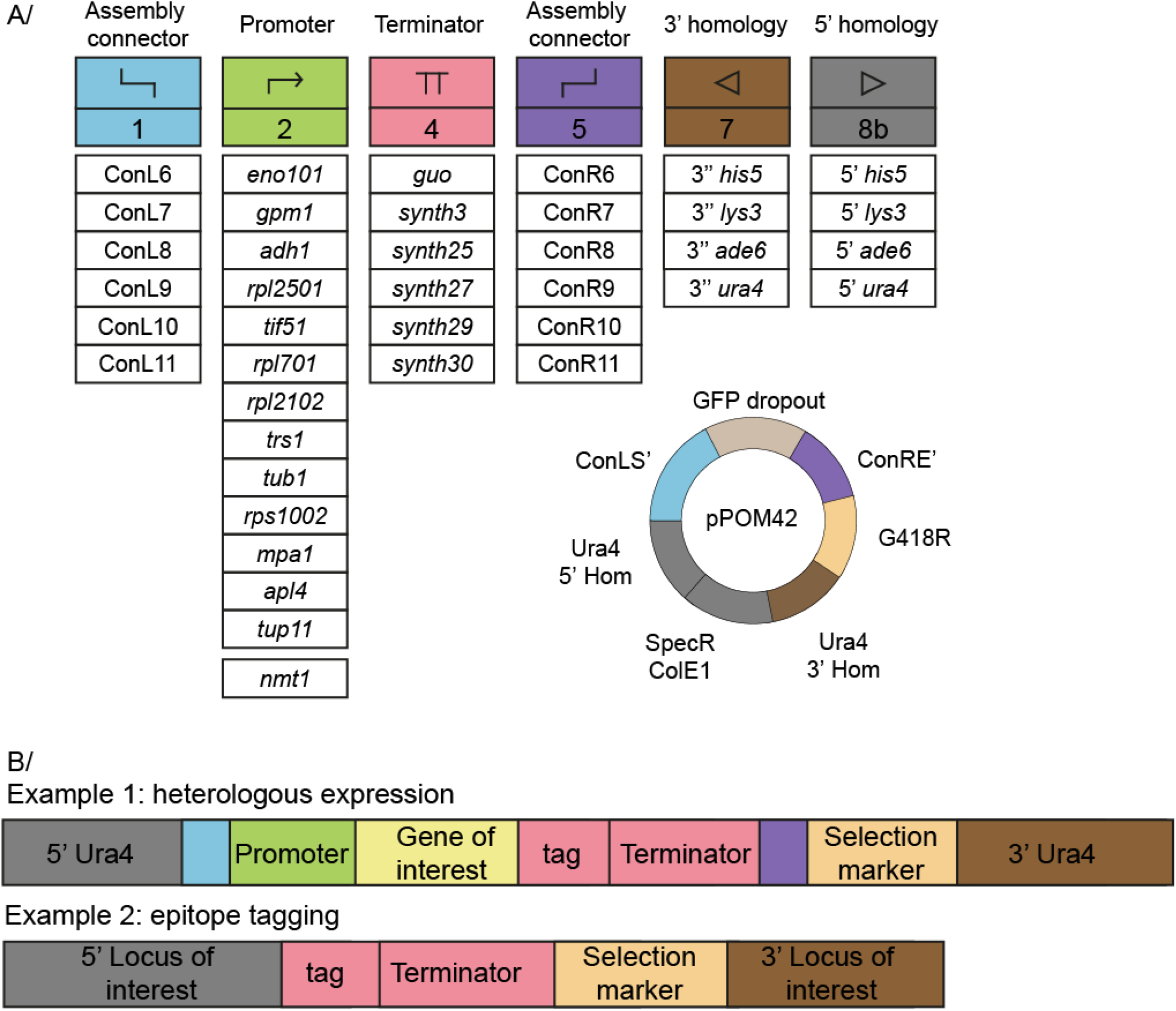
New parts provided by the POMBOX toolkit. (**A**) The toolkit provides six new assembly connector pairs (part type 1 and 5), fourteen characterized promoters (part type 2), six short synthetic terminator sequences (part type 4) and four pairs of homology arms (part type 7 and 8b), domesticated from Stable Integration Vectors.^20^ The POMBOX toolkit also included two backbone vectors for genomic integration such a pPOM042 for multi-gene assembly. (**B**) Two applications from POMBOX DNA parts. Example 1 highlights a tagged protein overexpression vector, and example 2 a vector for epitope tagging.

### New connectors for multi-gene plasmid assembly

Connector parts enable the generation of multi-gene plasmids from single transcriptional unit plasmids (e.g., a promoter, a coding sequence, and a terminator). Each pair of connectors features a BsmBI site and a unique overhang for assembling the transcriptional units in a systematic manner. The original YTK toolkit included six connector pairs for multi-gene assembly, but there is growing demand for larger pathways that can produce specialized metabolites in *S. cerevisiae*^31^, and the Golden Gate approach has been used to build assemblies of 52 fragments up to 40 kb long.^32^ To meet this demand, we designed six new connectors for building larger pathways. The connectors include a 143-bp concatenation of barcode sequences, a BsmBI recognition site, unique overhangs, and a 21-bp barcode scar, as recommended by the YTK. To avoid homology with *S. pombe* or *S. cerevisiae* genomes, we selected barcode sequences with no similarity to these genomes.^33^ To ensure that the overhang sequences of the new connectors are accurate, we used the NEBridge Ligase Fidelity platform to achieve assemblies with over 99% fidelity.^29, 30^ We named the six new pairs ConL5-11 and ConR5-11.

### A collection of promoters for protein expression

We characterized the levels of gene expression for thirteen constitutive promoters and one regulatable promoter. Three of the fourteen characterized promoters (2 constitutive, P*tub1* and P*tif51*, and 1 regulatable, P*nmt1*) are also used in the pDUAL2 vector series.^34^ To cover a wide range of gene expression with POMBOX, we selected eleven additional promoters. To do so, we used transcriptomic data generated by Thodberg *et al*. In this study, they analyzed the *S. pombe* transcriptome under two sets of standards culture conditions (EMM2 and Yeast Extract with Supplement (YES) media) and three sets of stress-inducing condition in *S. pombe* cell physiology (YES with 15 minutes at 39 °C, EMM2-with nitrogen starvation, and YES with 0.5 mM H2O2 for 15 min).^35^ Thus, promoters showing < 5% variability of transcript per million between the five sets of conditions we retained as constitutive. POMBOX selected promoters cover a range of expression levels (expressed as transcripts per million) from 120 to 300,000 (2,500-fold change, **Figure S2**). The promoters were amplified from the genome of *S. pombe* 972h-. They consist of the 5‘ UTR and the 1,000 bp downstream of the transcription start site. *Ptub1* (also named atb2) and P*tif51* were amplified from the pDUAL2 plasmid series. To evaluate the strength of each promoter, we cloned them upstream of a fluorescent reporter (*mRuby2* or *Venus*)^36, 37^ and measured fluorescence during exponential growth in YES or EMM2 using flow cytometry (**Figure 3, S3**). Each construct was made using the same terminator (T*eno1*) and integration locus (ura4).

**Figure 3:**
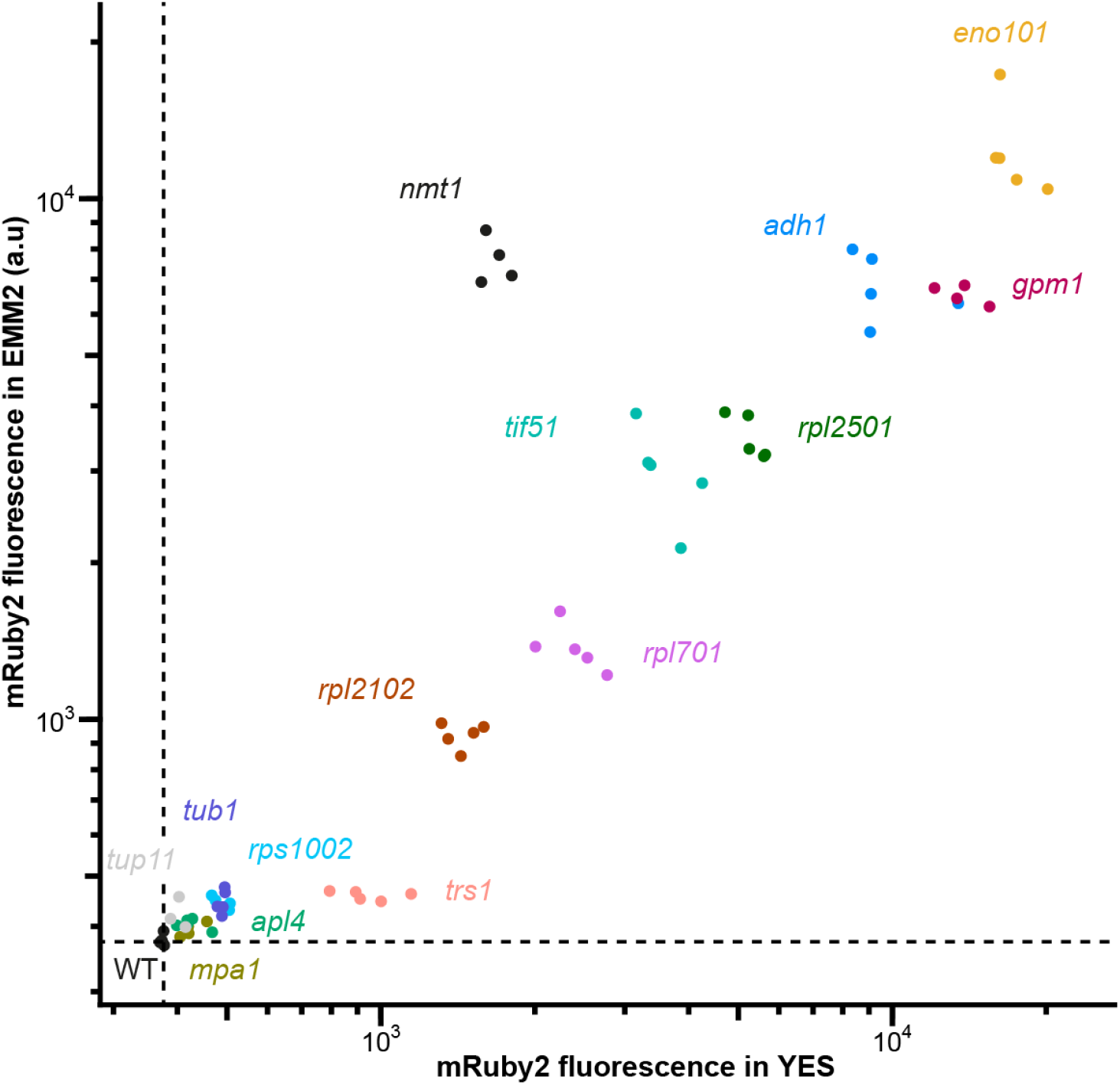
Strength of POMBOX promoters. The strength of fourteen promoters was quantified by flow cytometry by measuring the fluorescence emitted by the mRuby2 protein under the control of each promoter in the EMM2 or YES medium. WT is the background fluorescence from wild type *S. pombe* cells.

The promoters we propose cover 30 to 40-fold fluorescence intensity values and have been tested in two different media, YES and EMM2. These promoters are constitutive, with P*nmt1* as a regulatable promoter. For the constitutive promoters, the medium’s composition has no to little impact on the protein expression. Virtually all *S. pombe* native promoters are compatible with the toolkit and can extend the fourteen that we have provided. For interoperability between experiments and labs, we also present relative fluorescence values of the fourteen promoters normalized to P*adh1* fluorescent protein expression (**Table S9**).

### A collection of terminators for protein expression

Homologous recombination events after genomic integration, due to sequence homology, are undesirable in molecular or synthetic biology applications. The simplest and most effective way to minimize these events, if possible, is to use exogenous sequences that are not similar to genomic sequences. To achieve this, using sequences from evolutionarily distant organisms or synthetic sequences is a good practice. It has been shown that transcription terminators can be transferred from *S. cerevisiae* to *Pichia pastoris* while maintaining similar protein production capabilities, and that synthetic terminators designed and tested in *S. cerevisiae* also retain their properties in *P. pastoris*.^38^ This has also been shown to be the case empirically in *S. pombe*, since the T*adh1* terminator used in pDUAL vector series, as well as the T*cyc1* terminator used in the Stable Integration Vectors,^20^ are terminators from *S. cerevisiae*. We therefore verified that the six *S. cerevisiae* terminators present in the YTK, including T*adh1*, as well as six short synthetic terminators, can be used in *S. pombe*^39, 40^ and maintain protein production levels similar to *Tadh1*. To evaluate the protein production level of each terminator, we cloned it downstream of a fluorescent reporter (mRuby2) and measured fluorescence during exponential growth in EMM2 using flow cytometry (**Figure 4**). Each construct was made using the same promoter Padh1 and integration locus ura4.

**Figure 4:**
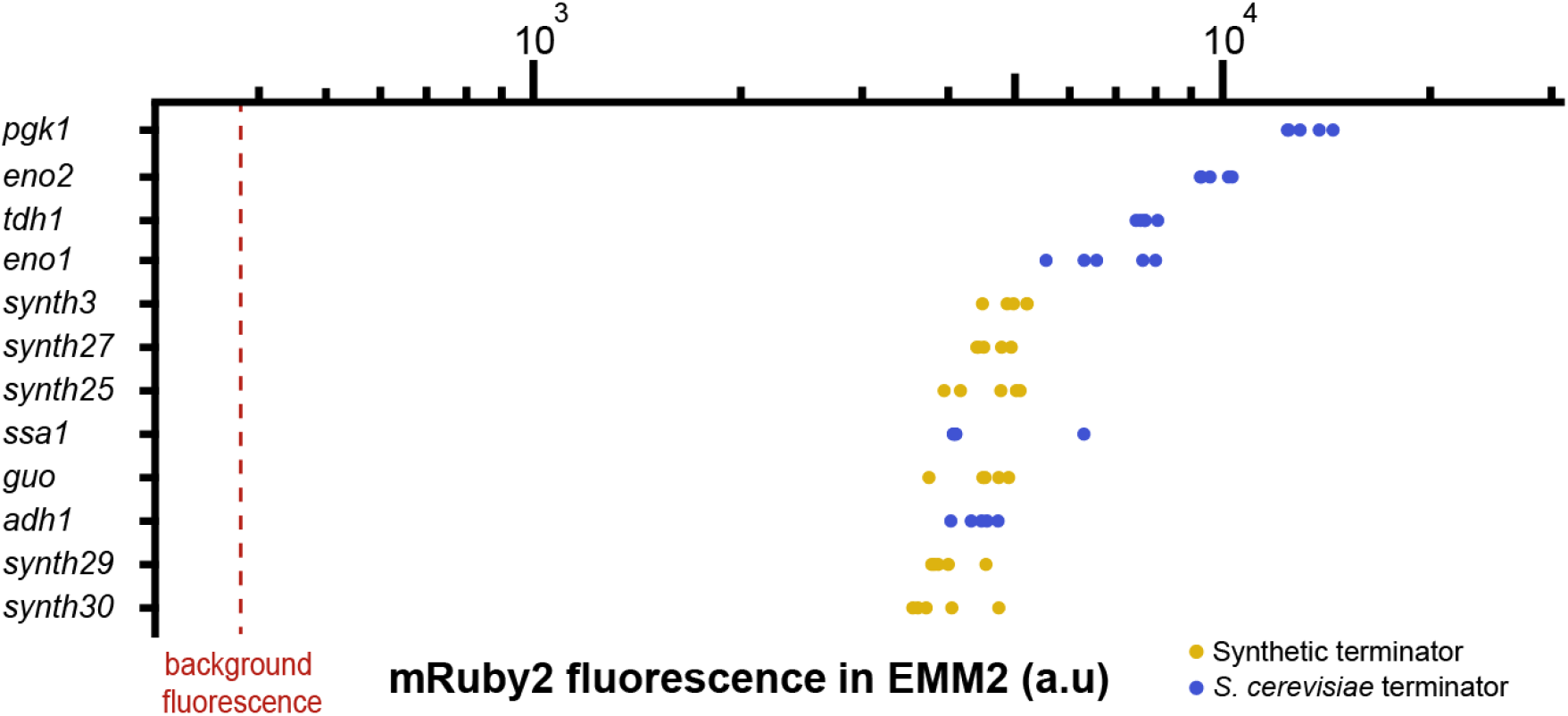
Strength of POMBOX terminators. The strength of six synthetic terminators and six *S. cerevisiae* terminators from the MoClo-YTK toolkit was quantified by flow cytometry by measuring the fluorescence emitted by the mRuby2 protein under regulation of each promoter in EMM2.

The expression difference we measured between the strongest T*pgk1* and weakest T*synth30* terminators was a 3.4-fold change. All terminators from *S. cerevisiae* provided stronger expression values than the commonly used T*adh1*. Overall, the synthetic terminator displayed lower expression values, similar to T*adh1* and T*ssa1* from *S. cerevisiae*. On the other hand, synthetic terminators provide a much smaller DNA sequence compared to terminators provided by the yeast toolkit. The synthetic terminators range from 50 to 80 bp whereas the YTK terminators are 230 bp long.

### S. pombe as a chassis for metabolic engineering

To emphasize the utility of *S. pombe* as a metabolic engineering chassis, we redirected three metabolic pathways (purine, mevalonate and amino acid) to build some precursors of specialized metabolites (**Figure 5**). Monomethylxanthine methyltransferase 1 (MXMT1) from *Coffea arabica* has been described to catalyze the methylation of xanthine, leading to the production of caffeine or theophylline.^41^ We expressed *Ca*MXMT1 under the regulation of P*adh1* and T*eno1*. After analysis of the ethyl acetate extract from *S. pombe* expressing *Ca*MXMT1 using LC-ESI-HRMS/MS, two new peaks with an *m/z* of 167.0563 (calculated for [C_6_H_6_N_4_O_2_ + H]^+^, −0.5 ppm) were found in the chromatogram (**Figure 5A**). It shows an effective production of methylxanthines from xanthine by *Ca*MXMT1, supported by MS/MS library matching (**Figure S4**) and an increase in retention time. Amorpha-4,11-diene synthase 1 (AMS1) from *Artemisia annua* catalyzes the first reaction toward the production of artemisinin, a sesquiterpenoid with antimalarial properties. We expressed *Aa*AMS1 under the regulation of P*adh1* and T*eno1*. After analysis of the ethyl acetate extract from *S. pombe* expressing *Aa*AMS1 using GC-EI-MS, a new peak of *m/z* 204.2 was found (**Figure 5B**). The MS spectrum of this compound was matched against the NIST library, identifying amorpha-4,11-diene as the best match (**Figure S5**). Phenyl ammonia lyase 2 (PAL2) from *Arabidopsis thaliana* catalyzes the transformation of phenylalanine into cinnamic acid, the first step of the biosynthesis of phenylpropanoids. We expressed *At*PAL2 under the regulation of P*ADH1* and T*eno1*. After analysis of the ethyl acetate extract from *S. pombe* expressing *At*PAL2 using LC-ESI-HRMS/MS, a new peak corresponding to cinnamic acid was found, confirmed by the analysis of a cinnamic acid standard with the same method (**Figure 5C**).

**Figure 5:**
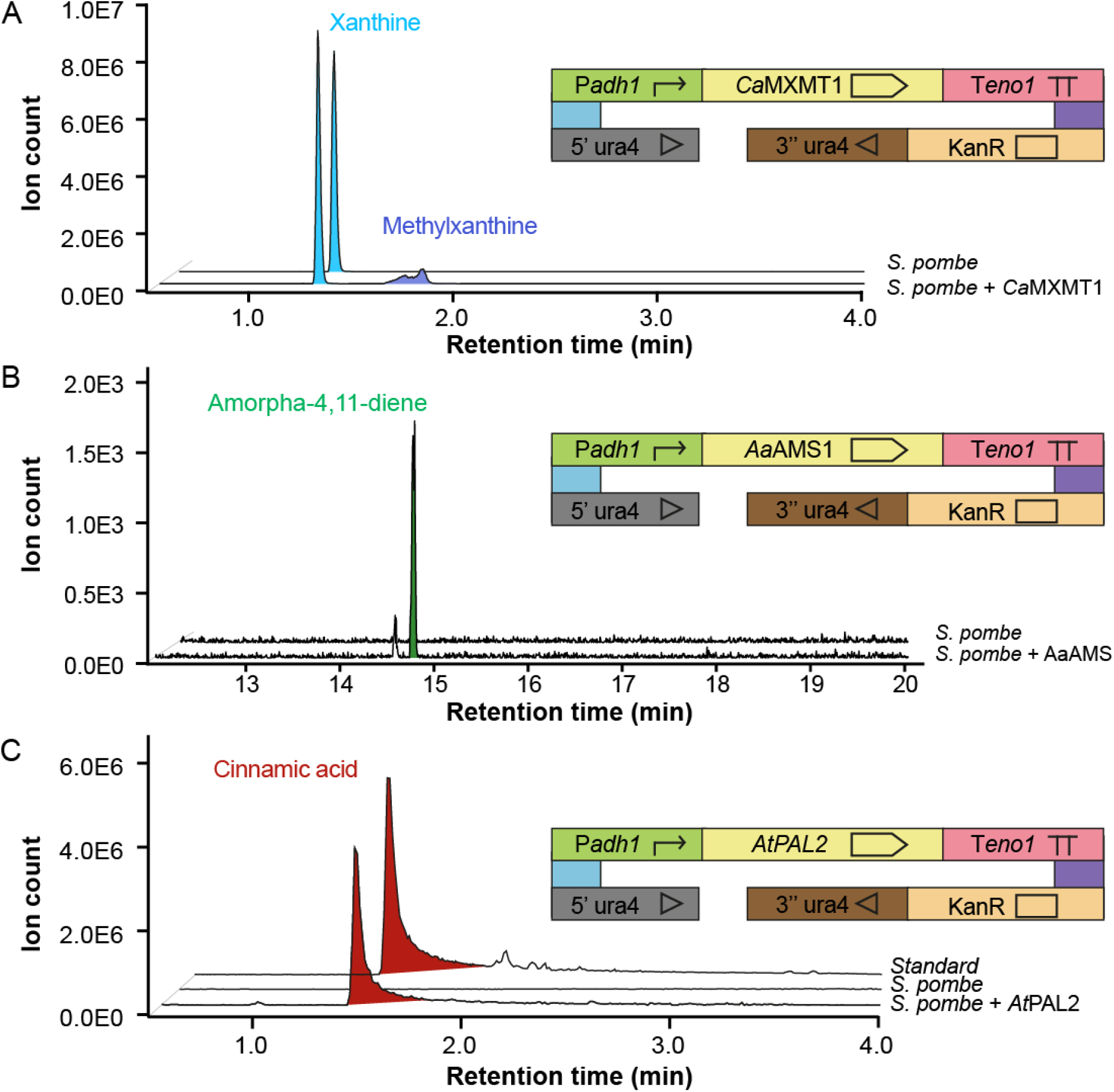
Production of metabolites in a single enzymatic step. (**A**) production of methylxanthine from xanthine (**B**) production of amorpha-4,11-diene from farnesyl pyrophosphate. (**C**) production of cinnamic acid from phenylalanine.

Thus, using POMBOX, we have successfully integrated into the *S. pombe* genome three different plant enzymes initiating specialized metabolites pathway and successfully producing the metabolites of interest.

### Genomic integration efficiency of large integration vectors in S. pombe

To test the genomic integration efficiency of large DNA sequences in *S. pombe*, we generated multi-gene plasmids using a combination of MoClo-YTK and POMBOX parts. Those plasmids included two to eight cassette plasmids (**Figure 6A**). Because we wanted to avoid lethal protein overexpression in this experiment, we chose *S. cerevisiae* promoters from the MoClo-YTK toolkit to avoid the expression of coding sequences in *S. pombe*. Purified plasmids were diluted to the exact same number of copies (10^−13^ mol per transformation) and were transformed into *S. pombe*. Integration vectors were targeted at the ura4 locus. DNA sequences up to 19 kb were successfully integrated into the *S. pombe* genome with a transformation efficiency range of 0.895–0.768 colony-forming units per 10^5^ cells. These values are on the same order of magnitude as the one reported in the study describing the stable integration vectors.^20^ We observed a slight reduction of the transformation efficiency depending on the size of the DNA sequence.

**Figure 6:**
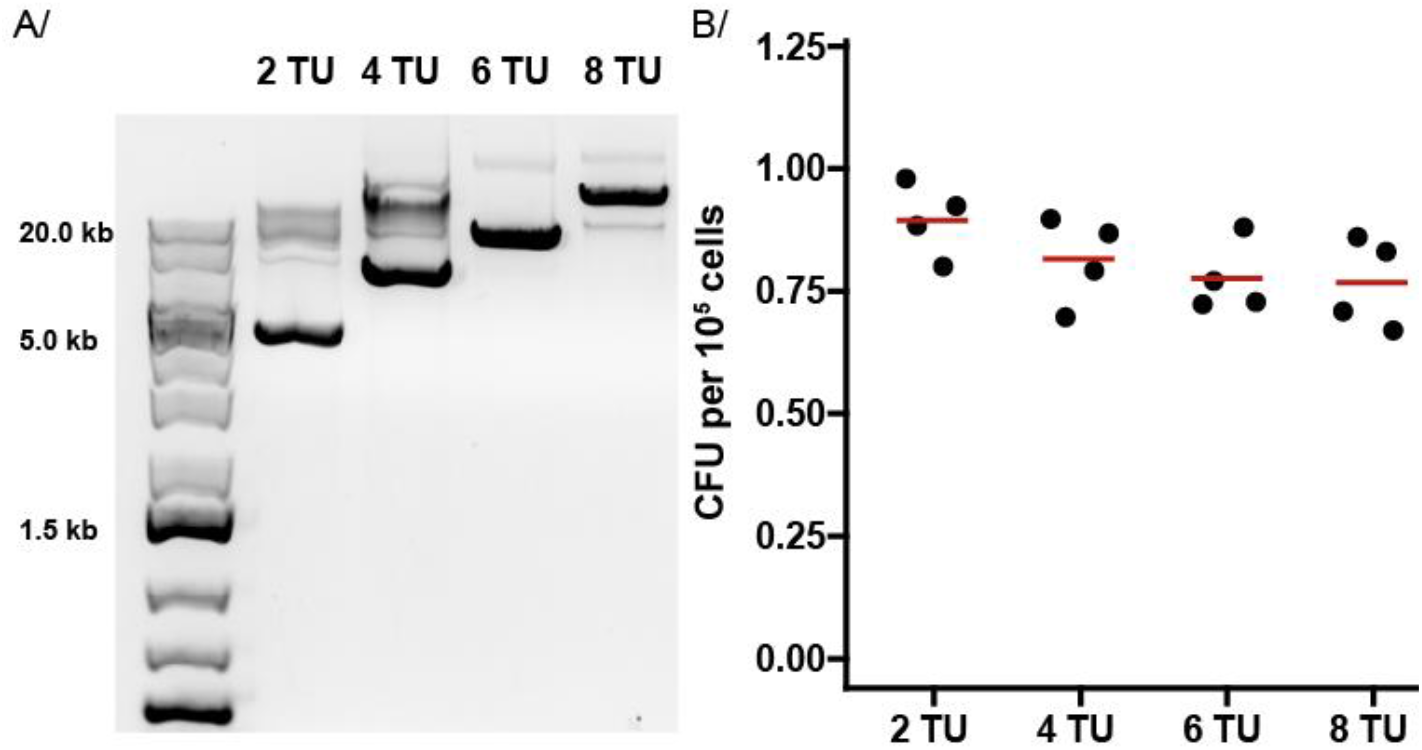
multi-gene integration vectors for S. pombe. (**A**) Plasmids from two to eight transcriptional units were generated, corresponding to a 4–19 kb sequence to be integrated into the *S. pombe* genome. (**B**) Transformation efficiency of *S. pombe* in function of integration vector size, from 4 to 19 kb. CFU: colony-forming units.

## Conclusions

To support the adoption of *S. pombe*, a model organism in molecular biology and cell physiology, as a chassis for synthetic biology, we developed POMBOX, a collection of parts compatible with the MoClo-YTK toolkit for hierarchical Golden Gate assembly. As a first step toward the use of *S. pombe* as a chassis organism in synthetic biology, we characterized fourteen promoters, six exogenous and six synthetic terminators, and expressed three plant enzymes that successfully produced precursors of high-value metabolites.

## Materials and Methods

### Growth Media, Strains

Chemicals used for media preparation were purchased from either Sigma–Aldrich (St. Louis, Missouri, United States), Duchefa Biochemie (Haarlem, Netherlands), Lach:ner (Neratovice, Czech Republic) or Penta chemicals (Prague, Czech Republic). Solvents for metabolic sample preparation, LC– and GC–MS were purchased from Fisher Chemical (Waltham, Massachusetts, United States) and were LC–MS grade. EMM2 medium was prepared according to Petersen & Russell^42^ with ammonium chloride, 5 g·L^−1^, potassium hydrogen phthalate 3 g·L^−1^, Na_2_HPO_4_, 2.2 g·L^−1^, glucose 20 g·L^−1^, salt stock 50×, vitamin stock 1,000× and mineral stock 10,000×. YES medium was prepared according to Petersen & Russell^42^ with 30 g·L^−1^ glucose, 5 g·L^−1^ yeast extract, 0.2 g·L^−1^ adenine, 0.2 g·L^−1^ uracil, 0.2 g·L^−1^ histidine, 0.2 g·L^−1^ leucine and 0.2 g·L^−1^ lysine. For solid media, 20 g·L^−1^ of agar was added.

*S. pombe* strains *972h-* and *975h+* ura4 were used as starting strains. *S. pombe* was used as a template for all DNA parts amplifications. *S. pombe* h+ 975 ura4 was used as a starting strain for genetic engineering. The complete list of strains generated for this study is available in Table S6.

DH10*β* electrocompetent *E. coli* cells were used for all molecular cloning experiments. Transformed cells were selected on Lysogeny Broth (LB) with the appropriate antibiotics (ampicillin, chloramphenicol, or spectinomycin). SOC was used for recovery after electroporation.

### Growth condition

For maintenance, *S. pombe* strains were cultivated in YES or EMM2 solid media at 28°C.

For general preculture, *S. pombe* strains were cultivated in YES or EMM2 at 28 °C, 200 RPM in an orbital shaker.

For experiments measuring promoter and terminator strength, *S. pombe* strains were cultivated in 1-mL volumes on 96-deep-well plates (CR1496, Enzyscreen, Hamburg, Germany, Netherlands) sealed with AeraSeal (Excel Scientific, Victorville, California), at 28 °C, 1,500 RPM on an Eppendorf ThermoMixer C (Eppendorf,Hamburg, Germany).

For metabolic pathway expression experiments, *S. pombe* strains were cultivated in EMM2 on 24- deep-well plates (CR1426, Enzyscreen, Hamburg, Germany, Netherlands) sealed with AeraSeal™ (Excel Scientific, Victorville, California), at 28 °C, 800 RPM on an Eppendorf ThermoMixer® C (Eppendorf, Hamburg, Germany).

For plasmid amplification, *E. coli* strains were cultivated in LB medium in 24-deep-well plates (CR1426, Enzyscreen, Hamburg, Germany, Netherlands) sealed with AeraSeal™ (Excel Scientific, Victorville, California), at 37 °C, 800 RPM on an Eppendorf ThermoMixer® C (Eppendorf, Hamburg, Germany).

### Plasmids

The list of all plasmids used in the study is available in Table S1-5.

All pPOM and pTP plasmids were generated using Golden Gate assembly based on the toolkit and overhangs of Lee et al.^21^

Part plasmids (Table S1, S2, S3) use pYTK001 as a backbone. DNA parts were either synthesized by TwistBioscience (South San Francisco, California, United States) as gene fragments (*Aa*AMS1, *Ca*MXMT1, ConL6-11, ConR6-11), Genscript (Piscataway, New Jersey, United States) subcloned in PUC57-BsaI-BsmBI-free (T*guo*, T*synth3*, T*synth25*, T*synth27*, T*synth29*, T*synth30*) or amplified from the *S. pombe* 972h- genome, pDUAL-FFH21 (P*tub1*) and pDUAL-FFH51 (P*tif51*) using overhang PCR. In some cases (P*nmt1*, P*adh1*, 5′-ura4, 5′-ade6, 3′-lys3, 5′-lys3, 3′-his5), a mutation was inserted using overlap extension PCR to remove BsmBI or BsaI restriction sites.

Cassette plasmids (Table S4) use pPOM041 or pYTK095 as a backbone. pPOM41 was used for direct integration of cassettes into the *S. pombe* genome and pYTK095 for multi-gene plasmid assembly. multi-gene plasmids (Table S5) use pPOM042 as a backbone.

Plasmid extraction was achieved from 2 mL of LB of an *E. coli* plasmid-harboring overnight culture, using a QIACube robot (Quiagen, Hilden, Germany) and the QIAprep Spin Miniprep Kit (27104, Quiagen), following the QIAprep miniprep protocol, the either “rapid” option for plasmids<10 kb or the “plasmid 10 kb or larger” option for plasmids >10 kb. Plasmids were eluted in 50 µL ddH_2_O.

### Polymerase chain reactions (PCR)

For the amplification of DNA parts from *S. pombe* or plasmids and overlap extension PCR, we used Phusion® High-Fidelity DNA Polymerase (M0530L, New England Biolabs, Ipswich, Massachusetts, United States) with the following conditions:

In 20 µL final, GC buffer 4 µL, 10 mM dNTPs 0.4 µL, 10 µM forward primer 1 µL, 10 µM reverse primer 1 µL, DNA template 1 µL, Phusion polymerase 0.2 µL, ddH_2_O12.6 µL. Reactions were conducted in a ProFlex™ 3 × 32-well PCR System thermocycler (Applied Biosystem, Waltham, Massachusetts, United States). Thermocycling conditions used the following template: 98 °C for 5 min as initial denaturation, for 35 cycles: 98 °C for 15 s, annealing temperature for 15 s, 72 °C for 30s·kb^−1^ and final extension 72 °C for 5 minutes. PCR products were separated on agarose gels (1 % w/v), 90 V, 60 min and purified using NucleoSpin® Gel and PCR Clean-up (Macherey-Nagel, Düren, Germany) before subsequent use.

For colony PCR and yeast genotyping, we used the Phire Green Hot Start II PCR Master Mix (Thermo Fisher Scientific, Applied Biosystem, Waltham, Massachusetts, United States) with the following conditions:

In 10 µL final, Phire Green Hot Start II PCR Master Mix 5 µL, 25 µM forward primer 0.2 µL, 25 µM reverse primer 0.2 µL, DNA template 4.6 µL. Reactions were conducted in a ProFlex™ 3 × 32-well PCR System thermocycler (Applied Biosystem, Waltham, Massachusetts, United States). Thermocycling conditions used the following template: 98 °C for 2 min as initial denaturation, for 30 cycles: 98 °C for 10 s, annealing temperature for 10 s, 72 °C for 10 s·kb^−1^ and a final extension 72 °C for 2 minutes. PCR products were separated on agarose gel (0.8 % w/v), 130 V, 30 min.

*E. coli* colonies were selected with a toothpick and spotted 4 times on selective media. The remaining bacteria were thus resuspended in 10 µL ddH_2_O and boiled for 10 minutes before being used as a DNA template.

Yeast genotyping was adapted from Lõoke *et al*.^43^, *S. pombe* colonies were selected with a toothpick and resuspended in 100 μL 200 mM LiOAc, 1 % SDS solution and boiled for 10 minutes. Then, 300 µL of EtOH 96 % were added and the solution was a vortex and centrifuge 15,000× *g* for 3 minutes. The supernatant was discarded and the pellet washed with 500 µL EtOH 70% before centrifugation 15,000× *g* for 1 min. The supernatant was discarded and the pellet dried for 1 minute at room temperature. The precipitated DNA was dissolved in 100 µL ddH_2_O and cell debris spun down at 15 000× *g* for 1 min.

### Golden Gate assembly reaction

Part plasmids were generated in 10 µL reaction volume. T4 ligase buffer 1 µL, T4 ligase 0.5 µL (M0202L, New England Biolabs, Ipswich, Massachusetts, United States), BsmBI-v2 0.5 µL (R0739L, New England Biolabs, Ipswich, Massachusetts, United States), pYTK001 0.5 µL, DNA part 0.5 µL (20 fm), ddH_2_O up to 10 µL. Reactions were conducted in a ProFlex™ 3 × 32-well PCR System thermocycler (Applied Biosystem, Waltham, Massachusetts, United States). Thermocycling conditions used the following template: for 25 cycles, 42 °C for 2 min, 16 °C for 2 min, then 60 °C for 30 min and 80 °C for 10 min.

Cassette plasmids were generated in 10 µL reaction volume. T4 ligase buffer 1 µL, T4 ligase 0.5 µL (M0202L, New England Biolabs, Ipswich, Massachusetts, United States), BsaI-HF®v2 0.5 µL (R3733L, New England Biolabs, Ipswich, Massachusetts, United States), DNA parts 0.5 µL (20 fm), ddH_2_O up to 10 µL. Reactions were conducted in a ProFlex™ 3 × 32-well PCR System thermocycler (Applied Biosystem, Waltham, Massachusetts, United States). Thermocycling conditions used the following template: for 25 cycles, 37 °C for 5 min, 16 °C for 5 min, then 60 °C for 30 min and 80 °C for 10 min.

For backbone plasmid assembly, thermocycling conditions were modified to end after a long ligation step: for 25 cycles, 42 °C for 5 min, 16 °C for 5 min, then 16 °C for 30 min.

multi-gene plasmids were generated in 10 µL reaction volume. T4 ligase buffer 1 µL, T4 ligase 0.5 µL (M0202L, New England Biolabs, Ipswich, Massachusetts, United States), BsmBI-v2 0.5 µL (R3733L, New England Biolabs, Ipswich, Massachusetts, United States), DNA parts 0.5 µL (20 fm), ddH_2_O up to 10 µL. Reactions were conducted in a ProFlex™ 3 × 32-well PCR System thermocycler (Applied Biosystem, Waltham, Massachusetts, United States). Thermocycling conditions used the following template: for 25 cycles, 42 °C for 5 min, 16 °C for 5 min, then 60 °C for 30 min and 80 °C for 10 min.

### *E. coli* transformation

Electroporation cuvettes were cooled down at 4°C for 30 min before each experiment. A 20 µL volume of a suspension containing electrocompetent *E. coli* cells was thawed at 4°C for 10 min before each experiment. Once thawed, 0.5 µL of plasmid was added to the cells with gentle mixing. The electroporator was set to 1700 V. For chloramphenicol, spectinomycin and kanamycin selective markers, cells recovered in 1 mL of SOC for 1 h at 37 °C, 200 RPM. Thus, cells were concentrated to 100 µL through centrifugation at 5000× *g* for 3 min and plated to their respective LB + selection marker. In the case of ampicillin, cells were resuspended into 100 µL of SOC after electroporation and directly plated in LB + ampicillin. The cells were then grown overnight at 37 °C.

### *S. pombe* transformation

Fission yeast strains were generated by the standard lithium acetate transformation protocol^44^ and selected using auxotrophy.

A total of 500–1,500 ng of plasmid were digested using NotI-HF® in 8 µL final volume. Cells were precultured overnight in the YES medium, 28 °C, 200 RPM. Then, cells were diluted to OD 0.1 and cultivated in the YES medium, 28 °C, 200 RPM until they reached OD 0.5. For 5 mL of cells at OD 0.5: the cells were pelleted, 2,500× *g*, 5 min and washed in 5 mL of sterile ddH_2_O. The cells were pelleted again at 2,500× *g* for 5 min and resuspended in 1 mL sterile ddH_2_O, and pelleted again at 16,000× *g* for 1 min and resuspended in 1 mL TE/LiAc. Then, cells were concentrated in 100 µL TE/LiAc and 8 µL of digested plasmid and 10 µL of Salmon sperm were added, mixed gently and incubated for 10 min at room temperature. Then, 260 µL of TE/LiAc/PEG (40 % w/v) were added and the cells were incubated for 1 hour at 30 °C. 43 µL of DMSO were added and the cells were heat-shocked at 42 °C for 5 min, centrifuged at 6,000× *g* for 1 min and washed in water before plating on the EMM2 medium. The cells were grown for 3– 5 days at 28 °C.

### Flow cytometry

Cells were maintained in solid EMM2 media and their preculture in 1 mL EMM2 media on 96-deep-well plates (CR1496, Enzyscreen, Hamburg, Germany, Netherlands) sealed with AeraSeal™ (Excel Scientific, Victorville, California), at 28 °C, 1,500 RPM on an Eppendorf ThermoMixer® C (Eppendorf, Hamburg, Germany) for 1 day. The cells were seeded at OD 0.05 in 1 mL EMM2 media in 96-deep-well plates and cultivated for 16 h at 28 °C, 1,500 RPM. The cells were then washed in PBS and diluted to OD1 in PBS before analysis by flow cytometry.

Flow cytometry experiments were performed on an CytoFLEX LX Flow Cytometer (Beckman Coulter, Brea, California, United States) The following channels were used: B525-FITC (525/40 nm filter) and Y610-mCherry (610/20 nm filter). At least 15,000 events were acquired from singlet-gated populations using FSC-A/SSC-A.

Data were processed using the FlowJo software package with the following gating strategy: (1) the main cellular population was selected using forward and side scatter to exclude cell aggregates and debris, (2) doublets were excluded from analysis by plotting FSC-A versus FSC-H and gating along the diagonal. Number of cells and population mean were extracted to a .csv file. The data were then analyzed using the ggplot2 package in R. Each set of condition was tested five times in independent experiments.

### Metabolite extraction

Cells were maintained in solid EMM2 media and precultured in 1 mL EMM2 media on 96-deep-well plates (CR1496, Enzyscreen, Hamburg, Germany, Netherlands) sealed with AeraSeal™ (Excel Scientific, Victorville, California), at 28 °C, 1,500 RPM on an Eppendorf ThermoMixer® C (Eppendorf, Hamburg, Germany) for 1 day. Cells were seeded at OD 0.05 in 2 mL EMM2 media in EMM2 on 24-deep-well plates (CR1426, Enzyscreen, Hamburg, Germany, Netherlands) sealed with AeraSeal™ (Excel Scientific, Victorville, California), at 28 °C, 300 RPM for 5 days.

Then, 1 mL of ethyl acetate (E196-4, Fisher Scientific) was added to the culture medium and mixed for 1 hour at 28 °C, 250 RPM. Subsequently, 1 mL of ethyl acetate was added and thoroughly mixed by pipetting. Sample was collected in a 2 mL round bottom tube (Eppendorf, Hamburg, Germany) centrifuged for 5 mins, 14,100× *g*, and the organic phase was carefully collected and dried under N_2_ flow.

### LC–MS/MS

For LC**–**MS/MS analysis, samples were resuspended to 1 mg·mL^−1^ in an ACN:H_2_O (50:50) mixture. LC**–**MS analyses were performed using a Vanquish Flex UHPLC System interfaced to an Orbitrap ID-X Tribrid mass spectrometer, equipped with heated electrospray ionization (H-ESI). The LC conditions were as follows: column, Waters BEH (Ethylene-Bridged Hybrid) C18 50 × 2.1 mm, 1.7 μm; mobile phase, (A) water with 0.1 % formic acid; (B) acetonitrile with 0.1% formic acid; flow rate, 350 μL·min^−1^; column oven temperature, 40 °C, injection volume, 1 μL, linear gradient of 5 to 100 % B over 5 min and isocratic at 100 % B for 2 min. Electrospray ionization was achieved in positive mode and mass spectrometer parameters were as follows: ion transfer tube temperature, 325 °C, auxiliary gas flow rate 10 L·min^−1^, vaporizer temperature 350 °C; sheath gas flow rate, 50 L·min^−1^; capillary voltage, 3,000 V, MS resolution 60,000, quadrupole isolation, scan range from *m/z* 100–1,000, RF Lens 45 %, maximum injection time 118 ms. The data-dependent MS/MS events were acquired for the most intense ions from MS scan for a cycle time of 0.6 s, above a threshold of 1.0E5 intensity threshold, with a dynamic exclusion list of 2 s, including the isotopes. Selected precursor ions were fragmented with a fixed normalized HCD collision energy of 35 %, an isolation window of *m/z* 0.8 and a resolution of 15,000 with a maximum injection time of 80 ms.

### GC–MS

For GC–MS analysis, 1 mg of samples were resuspended into 100 µL ethyl acetate.

GC–MS analyses were performed using a 7890A gas chromatograph coupled with a 5975C mass spectrometer, equipped with electron ionization (EI) and quadrupole analyzer (Agilent Technologies, Santa Clara, CA, USA). The samples (1 μl) were injected to split/splitless inlet in split mode (split ratio 10:1). The injector temperature was 250 °C. A DB-1ms fused silica capillary column (30 m × 250 μm; film thickness of 0.25 μm, J&W Scientific) was used for separation. The carrier gas was helium at a constant flow rate of 1.0 ml·min^−1^. The temperature program was: 40 °C (1 min), then 5 °C·min^−1^ to 100 °C, followed by 15 °C·min^−1^ to 230 °C. The temperatures of the transfer line, ion source and quadrupole were 320 °C, 230 °C and 150 °C, respectively. EI spectra (70 eV) were recorded from *m/z* 25 to 500.

### Data analysis

LC–MS/MS .raw data files were directly imported into MZmine 3.4.16.^45^ Extracted ion chromatograms for compounds of interest were generated using the *Raw data overview* module and exported as .pdf.

GC–MS .dx files were analyzed using OpenLab CDS 2.4. Extracted ion chromatograms for compounds of interest were exported as .csv files and plotted using the ggplot2 package in R.

Figures were generated using Rstudio and the following packages: ggplot2, gridExtra, patchwork, dplyr, forcats, ggthemes, ggprism, DescTools, tidyverse and scales. Adobe Illustrator CS6 was used for their graphic adjustment.

## ASSOCIATED CONTENT

Supporting Information

Supporting Information is available free of charge at […].

Tables S1–S5, lists of plasmids, Table S6, list of strains Tables S7,S8, lists of primers, Table S9, relative fluorescence to Padh1, Figure S1, MoClo-YTK parts that can be used with *S. pombe*, Figure S2, Promoter strength in transcripts per million, Figure S3, Strength of POMBOX promoters with Venus, Figure S4, MS/MS Spectral comparison of methylxanthine, Figure S5, EI–MS spectrum comparison of amorpha-4,11- diene.

## AUTHOR INFORMATION

Corresponding Author

Tomáš Pluskal – Institute of Organic Chemistry and Biochemistry of the Czech Academy of Sciences, Prague, Czech Republic

## Author Contributions

T.H. conceptualization, methodology, investigation, validation, writing, revisions.

H.S. investigation and validation of integration vectors.

B.E. investigation for promoters.

M.P. resources, writing, revisions.

T.P. conceptualization, resources, funding, project administration, supervision, writing, revisions.

## Notes

The authors declare no competing financial interest.

## ACKNOWLEDGMENTS

T.H. is supported by the IOCB Fellowship program. T.P. is supported by the Czech Science Foundation Grant 21-11563M and by the European Union’s Horizon 2020 research and innovation program under Marie Skłodowska-Curie grant agreement 891397. We thank Mitsuhiro Yanagida for providing the *S. pombe* strains and Fred Rooks for editing the manuscript.

## Supplementary Information

**Figure S1:**
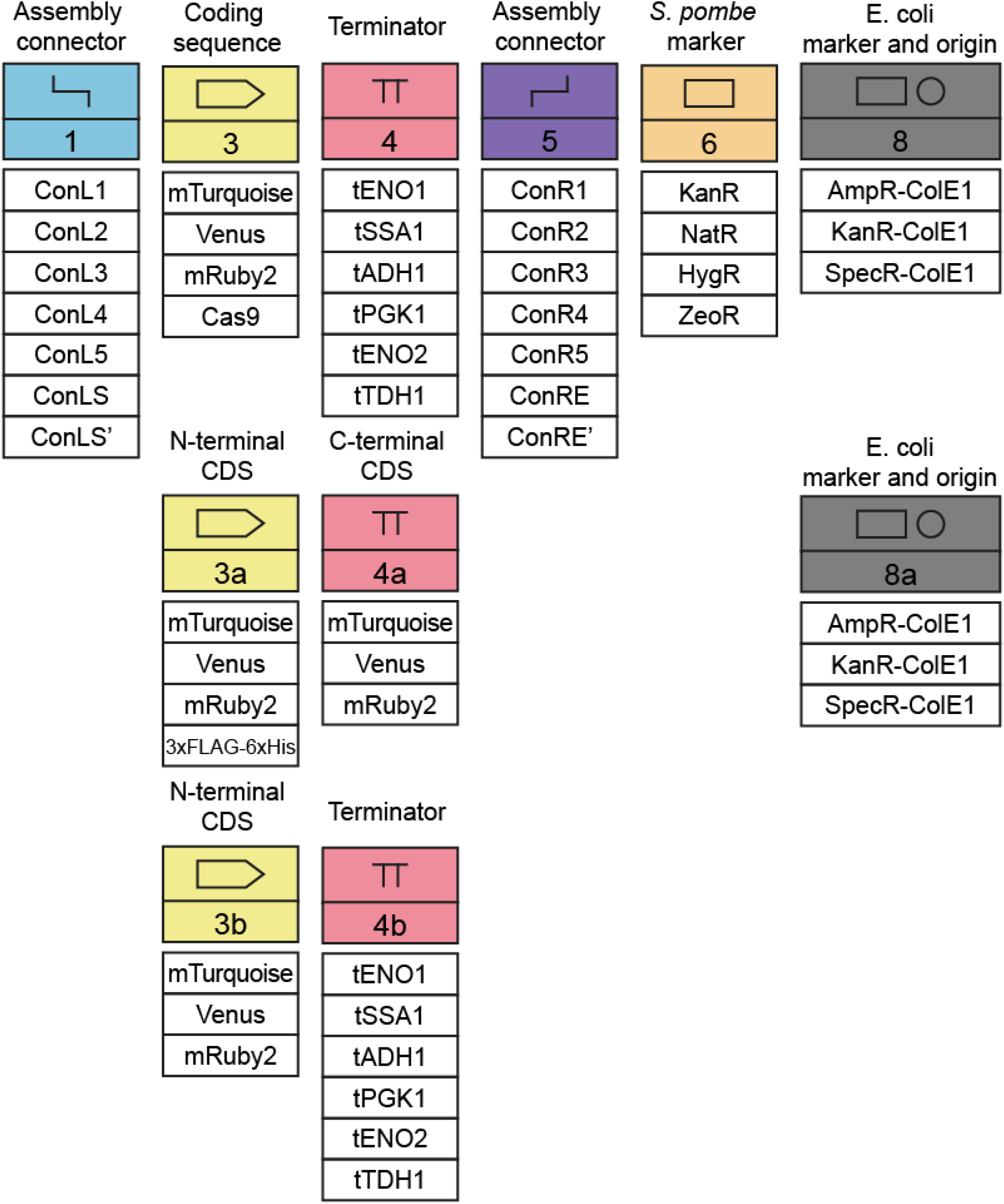
MoClo-YTK parts that can be used with *S. pombe*.

**Figure S2:**
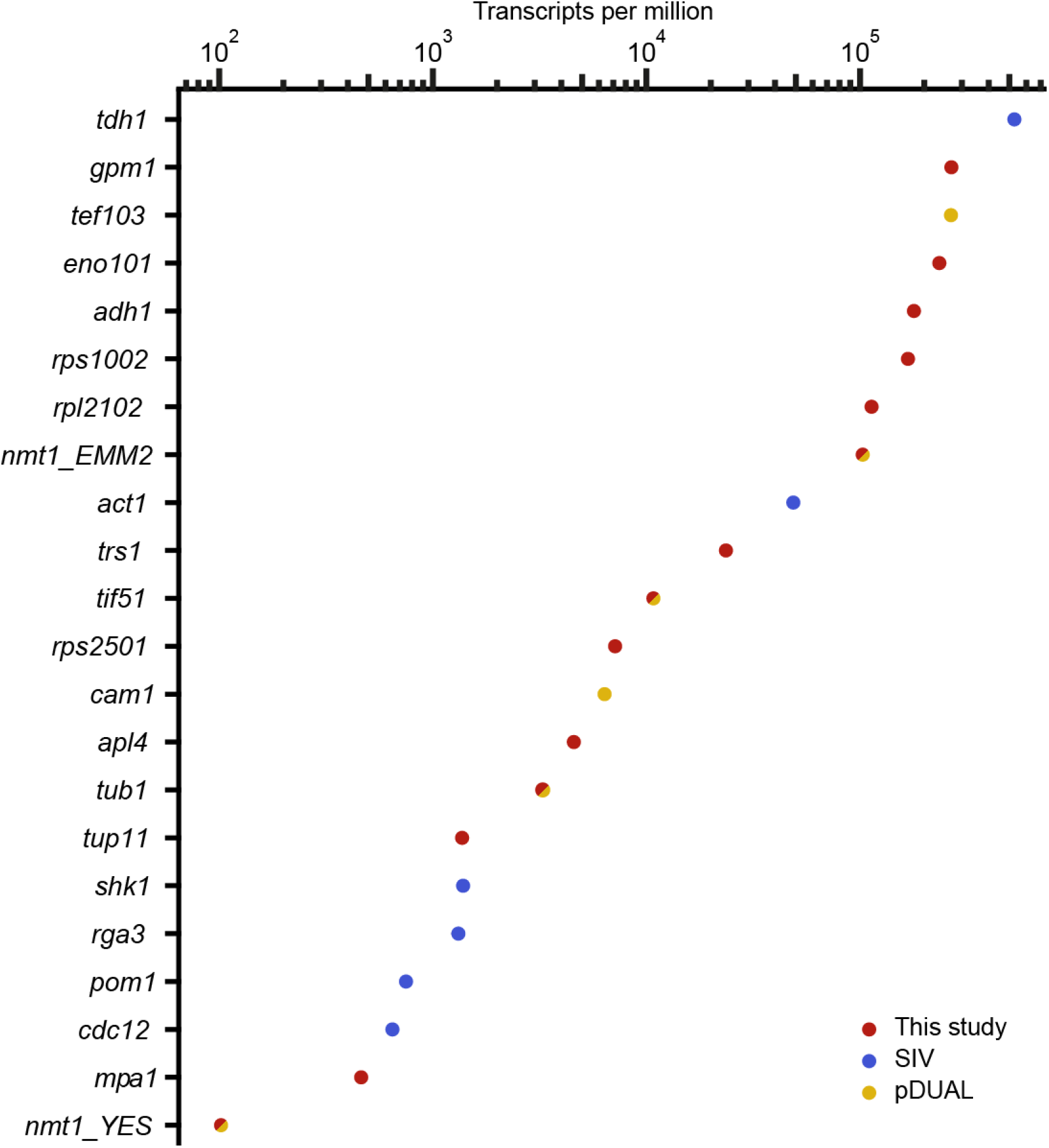
Promoters strength in transcript per million (TPM), measured by Thodberg et al.^35^ Red, selected for this study, blue, from Stable Integration Vectors tested by Vjestica et al.,^20^ yellow, in the pDUAL2 series.

**Figure S3:**
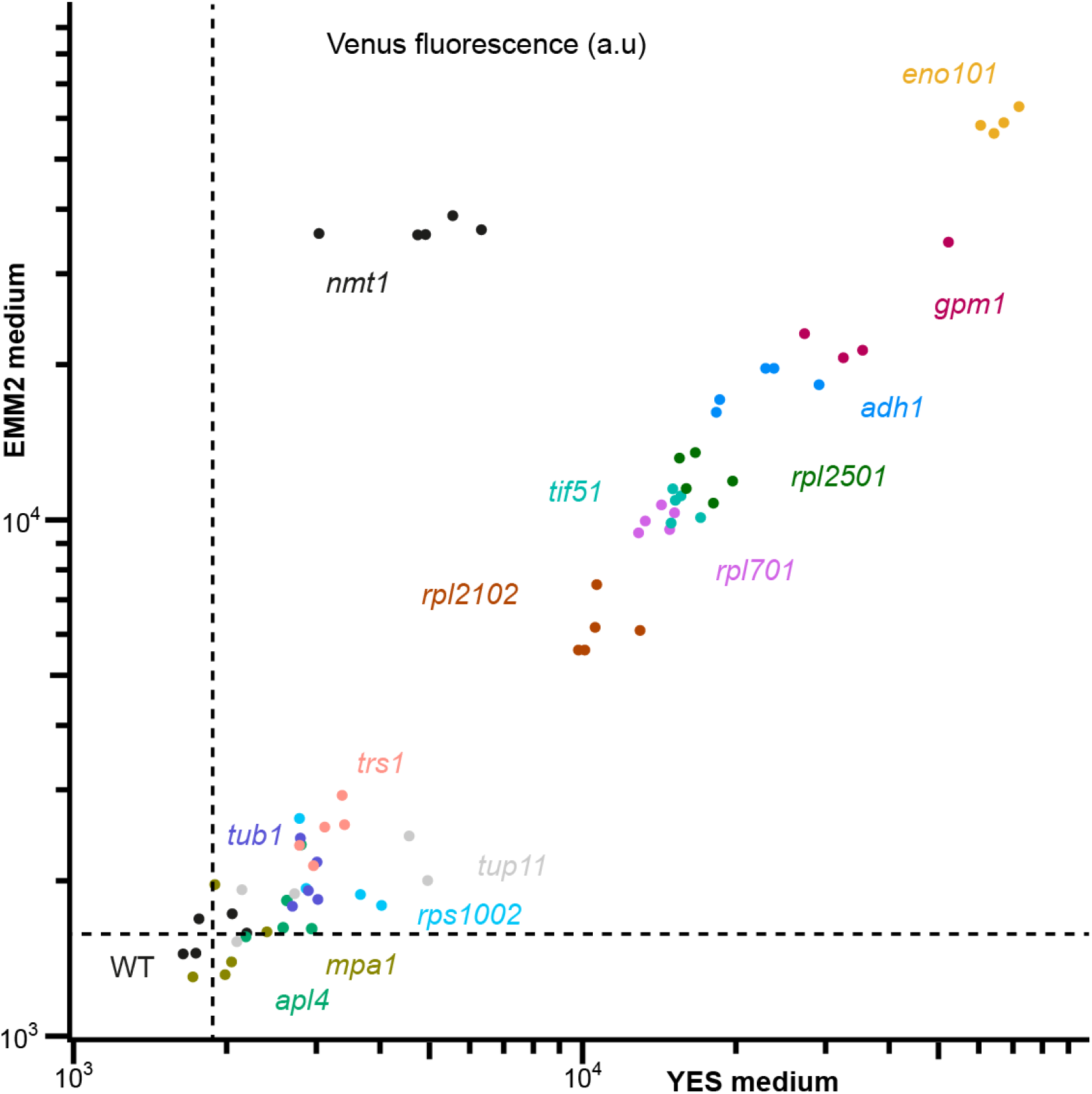
Strength of the POMBOX promoters. The strength of 14 promoters was quantified by flow cytometry by measuring the fluorescence emitted by the Venus protein under the control of each promoter in EMM2 or YES media. WT is the background fluorescence from *S. pombe* cells not expressing any fluorescent protein. P*nmt1* is a regulatable promoter repressed by thiamine, which is present in YES media.

**Figure S4:**
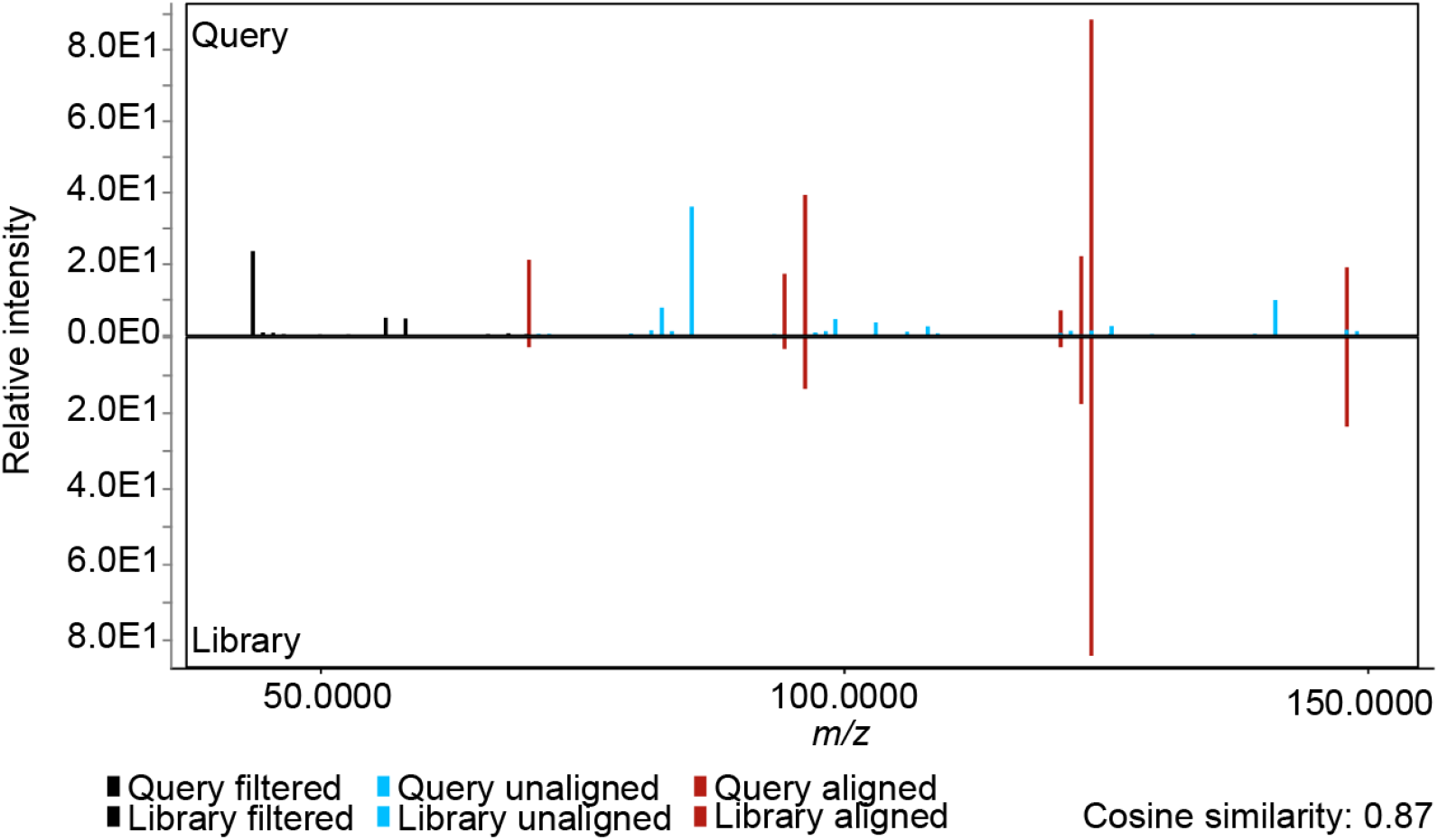
MS/MS Spectral comparison of methylxanthine produced by *S. pombe*_CaMXMT1 with 3- methylxanthine library match.

**Figure S5:**
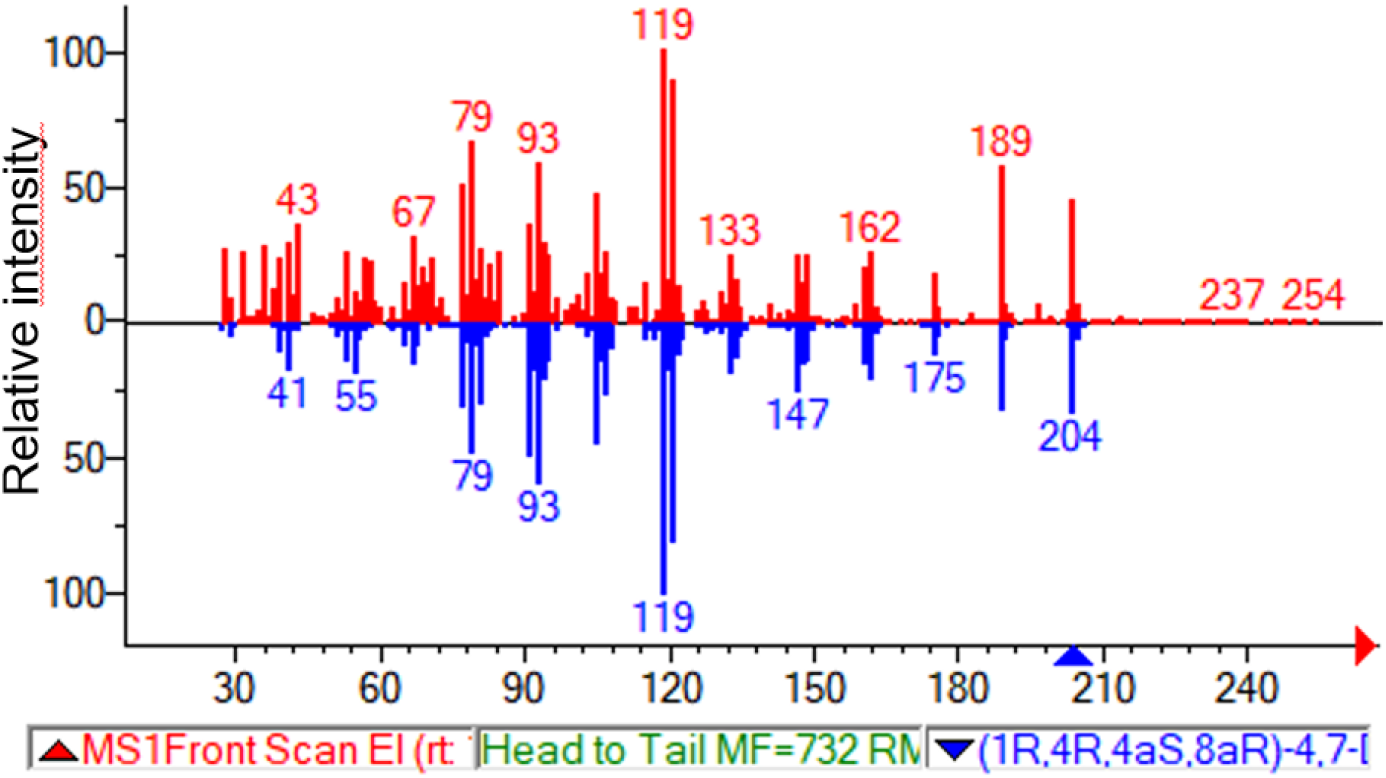
EI–MS spectrum comparison of amorpha-4,11-diene produced by *S. pombe*_AaAMS1 and amorpha-4,11-diene from NIST EI library.

### Tutorial on how to use POMBOX

To support the use of POMBOX, we present two illustrative examples of how to work with Golden Gate assembly and design new parts for molecular biology experiments.

*Example 1* is the most straightforward usage of POMBOX. It showcases a method of tagged protein overexpression targeting a classical prototrophy locus. It shows how to make a DNA sequence of interest compatible with POMBOX grammar and describes the experimental workflow.

*Example 1* illustrates how to domesticate DNA parts for use in Golden Gate assembly.

*Example 2* mimics the usage of pFA plasmids for epitope tagging. It aims to illustrate the versatility and modularity of Golden Gate assembly in molecular biology applications with more advanced design of sequences and choices of overhangs.

*Example 2* illustrates the versatility of Golden Gate assembly and how to tweak the overhang grammar for any molecular biology application.

For an in-depth understanding of the design of part types, we recommend reading the supplementary information associated with the MoClo-YTK article.

*Example 1: heterologous expression of a tagged protein in S. pombe*.

For this example, we will consider Cox4 and Cox5, which are two proteins whose structure has recently been resolved using epitope tagging in *S. pombe*^46^. Our goal is to overproduce proteins independently and to purify them with the help of the 6XHis tag.

Parts already proposed in POMBOX and MoClo-YTK.

With POMBOX and MoClo-YTK, some parts are already available and ready to use:

- **pPOM041** is the backbone plasmid for genomic integration. It holds parts 1, 5, 6, 7 and 8. It targets the Ura4 locus and brings KanR as a selection marker. pPOM041 also provides a GFP drop-out system for green-white colony screening.

- **pPOM013**: with P*eno101* is selected as part type 2. It is the strongest promoter available.

- **pYTK060**: with the 3XFLAG-6XHis tag

- **pYTK064**: with T*pgk1* is selected as part type 4b (terminators compatible with C-ter tags).

New DNA parts have to be generated.

For this specific study, two coding sequences (part type 3) have to be generated. Those parts are the coding sequence for the Cox4 and Cox5 proteins. There are two methods to obtain those parts:

A/ The coding sequences are synthesized as gene fragments.

B/ The sequences are obtained by amplification from the source organism.

Whatever the chosen approach, both sequences have to respect some core rules to be compatible with POMBOX and MoClo-YTK.

- The DNA sequence should be free of BsaI, BsmBI and NotI recognition sites.

- The overhangs should be part type 3 overhangs. See the list of overhangs at the end of the tutorial section for the exhaustive list.

For method A, gene fragment synthesis, we recommend designing sequences in the following way:

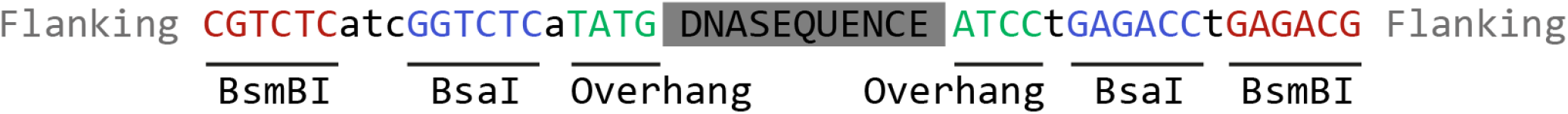

The sequence should include the coding sequence of interest (DNASEQUENCE, in this case Cox4 or Cox5), the **type 3 part overhangs**, the **BsaI recognition site** and the **BsmBI recognition site**. We also recommend adding some **flanking** nucleotides to facilitate the binding of the restriction enzymes.

Note that type 3 part overhangs already include the **start codon**.

For method B, amplification of the sequence, we recommend proceeding as follows:

- 1/ Check for any BsaI, BsmBI and NotI restriction sites in your sequence of interest. In the case of Cox4, there are none.

In the case of Cox5, there is one at position 376–381.

- 2a/ For the Cox4 sequence: DNA can be amplified by overhang PCR directly from the *S. pombe* genome. The set of PCR primers can be the following:

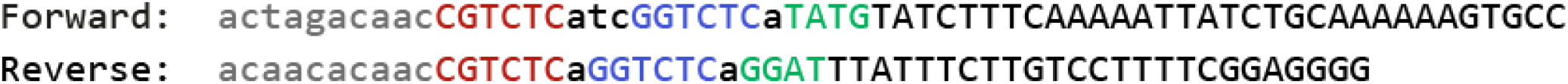

Note: We recommend using touch-down PCR and gel clean-up to obtain the DNA fragment of interest.

- 2b/ For the Cox5 sequence: DNA can be amplified directly from the genome using overhang and overlap PCR. The two primers can be the following:

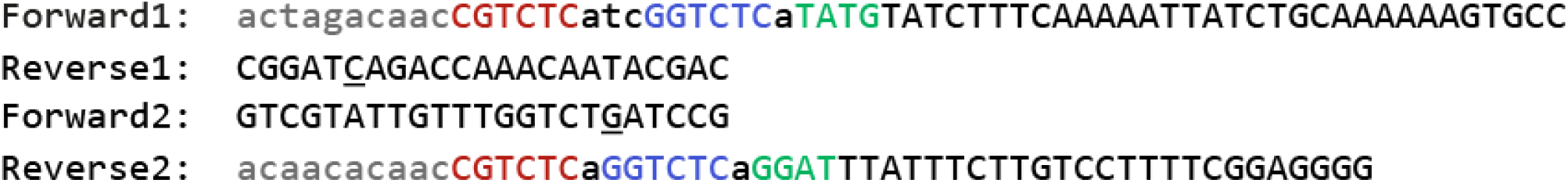

Note: The mutation is <u>underlined</u>. We recommend using touch-down PCR and gel clean-up to obtain the DNA fragment of interest.

- 3/ Store the DNA fragment in pYTK001. It uses BsmBI Golden Gate assembly, and pYTK001 possesses a GFP drop-out for green/white screen of transformant. If the DNA fragment was synthesized, we perform BsmBI Golden Gate assembly with a molar ratio of 1:1 of pYTK001 and the DNA fragment, 20 fm. If the DNA fragment was obtained by amplification, we perform BsmBI Golden Gate assembly with 20 fm of pYTK001 and 1 µL of the DNA fragment from gel clean up. We then use 0.5 µL of the Golden Gate reaction mix for 20 µL of competent cells.

- 4/ Perform the BsaI Golden Gate assembly to generate the integration vector of interest. We use a 20 fm equimolar ratio of plasmids holding the DNA parts of interest. Here: pPOM041 (backbone), pPOM013 (promoter), pCOX (coding sequence), pYTK060 (tag) and pYTK064 (terminator). We then use 0.5 µL of the Golden Gate reaction mix for 20 µL of competent cells. pPOM041 uses AmpR as a selection marker, so *E. coli* cells can directly be plated on the selection media, without phenotype expression. pPOM041 also provides a GFP drop-out system, for green-white colony screening.

- 5/ To integrate the DNA sequence of interest, we digest 0.5–5 µg of the resulting plasmid with NotI and use it with the LiAc/PEG transformation procedure.

*Example 2: Epitope tagging of a protein in the S. pombe genome*.

This example replicates the experimental set-up of Moe et al, for the protein Cox4.^46^ Our goal is to tag the C-ter of both Cox4 and Cox5 with TEV-6xGly-2xStrep.

### Already proposed parts for use with POMBOX and MoClo-YTK

For use with POMBOX and MoClo-YTK, some parts are already available and ready to use:

- **pYTK089** is the backbone plasmid. It holds the *E. coli* origin of replication and an ampicillin resistance marker. It also bears a RFP drop-out system for red/white colony screening.

- **pYTK063**: with T*adh1* is selected as part type 4b. (terminator after Cter tag)

- **pYTK065:** with ConR1 as part type 5

- **pYTK077**: with KanR selection marker as part type 6.

### New DNA parts have to be generated

The following parts are lacking in POMBOX and MoClo-YTK and have to be generated for this application:

- TEV-6xGly-2xStrep tag.

- 5′ Cox4

- 3′ Cox4

As in *example 1*, DNA sequences can be either synthesized as gene fragments or amplified from a template.

In both cases the key part of the design is related to the design of the overhangs.

TEV-6xGly-2xStrep tag is a regular part type 4a and therefore the overhangs can be as follows:

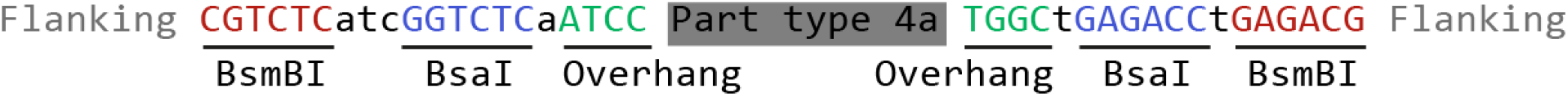

Cox4 3’ is a regular part type 7, so the overhangs can be as follows:

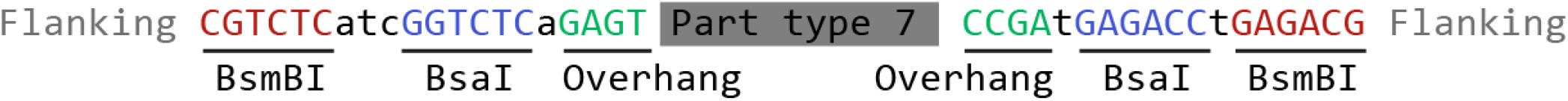

Cox4 5′ is a hybrid part, containing the 5′ overhang from part type 8b, and the 3′ overhang from part type 3. This way the only sequences that are inserted into the *S. pombe* genome are the TEV-6xGly-2xStrep tag, T*adh1*, ConnectorR1 and KanR.

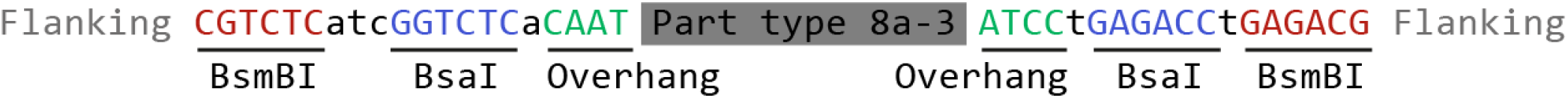

- 3/ Store the DNA fragment in pYTK001. It uses BsmBI Golden Gate assembly, and pYTK001 possesses a GFP drop-out for green/white screen of transformant. If the DNA fragment was synthesized, we perform BsmBI Golden Gate assembly with a molar ratio of 1:1 of pYTK001 and the DNA fragment, 20 fm. If the DNA fragment was obtained by amplification, we perform BsmBI Golden Gate assembly with 20 fm of pYTK001 and 1 µL of the DNA fragment from gel clean up. We then use 0.5 µL of the Golden Gate reaction mix for 20 µL of competent cells.

- 4/ Perform the BsaI Golden Gate assembly to generate the integration vector of interest. We use a 20 fm equimolar ratio of plasmids holding the DNA parts of interest. Here: pYTK089 (backbone), p5COX (5′ homology of Cox4), p3COX (3′ homology of Cox4), pYTK063 (terminator), pTEV-6xGly-2xStrep (tag), pYTK065 (connector), pYTK077 (selection marker).

We then use 0.5 µL of the Golden Gate reaction mix for 20 µL of competent cells. pYTK089 uses AmpR as a selection marker, so *E. coli* cells can be directly plated on the selection media, without phenotype expression. pPOM041 also provides a GFP drop-out system for green-white colony screening.

- 5/ To integrate the DNA sequence of interest, we digest 0.5–5 µg of the resulting plasmid with NotI and use it with the LiAc/PEG transformation procedure.

List of overhangs:

BsmBI for pYTK001 ligation:

ATCG

BsaI for cassette plasmid assembly:

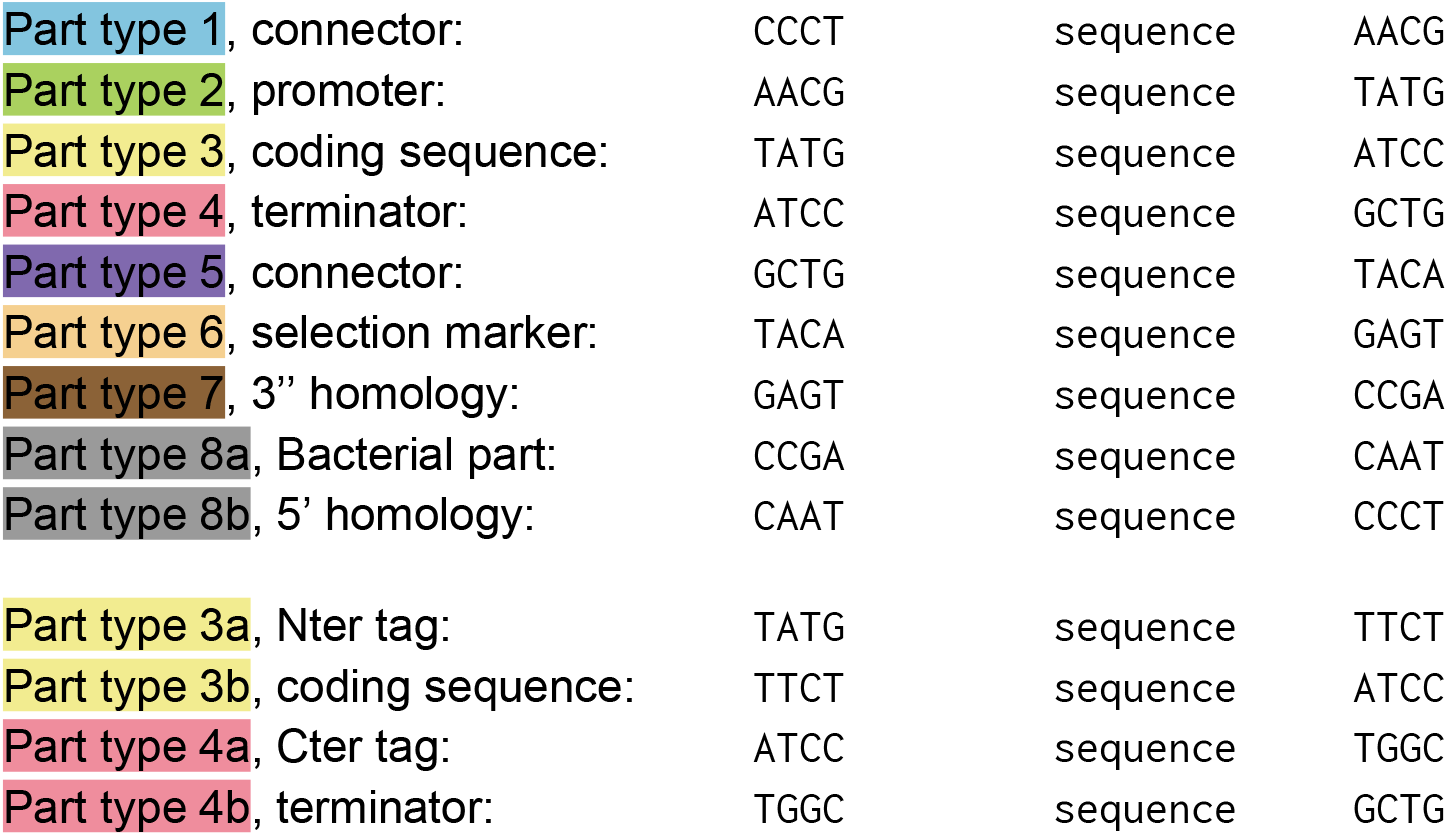

### List of sequences

Overhang PCR primers were designed based on the Genome assembly [ASM294v2](https://www.ncbi.nlm.nih.gov/data-hub/genome/GCF_000002945.1/) or plasmid sequences. The following overhang were added to primers: Forward overhang 5′ actagacaacCGTCTCatcGGTCTCaNNNN 3′

Reverse overhang 5′ acaacacaacCGTCTCaGGTCTCaNNNN 3′

The PCR overhangs include Golden Gate overhangs related to the part type (NNNN), BsaI and BsmBI restriction sites.

**Table S1:**
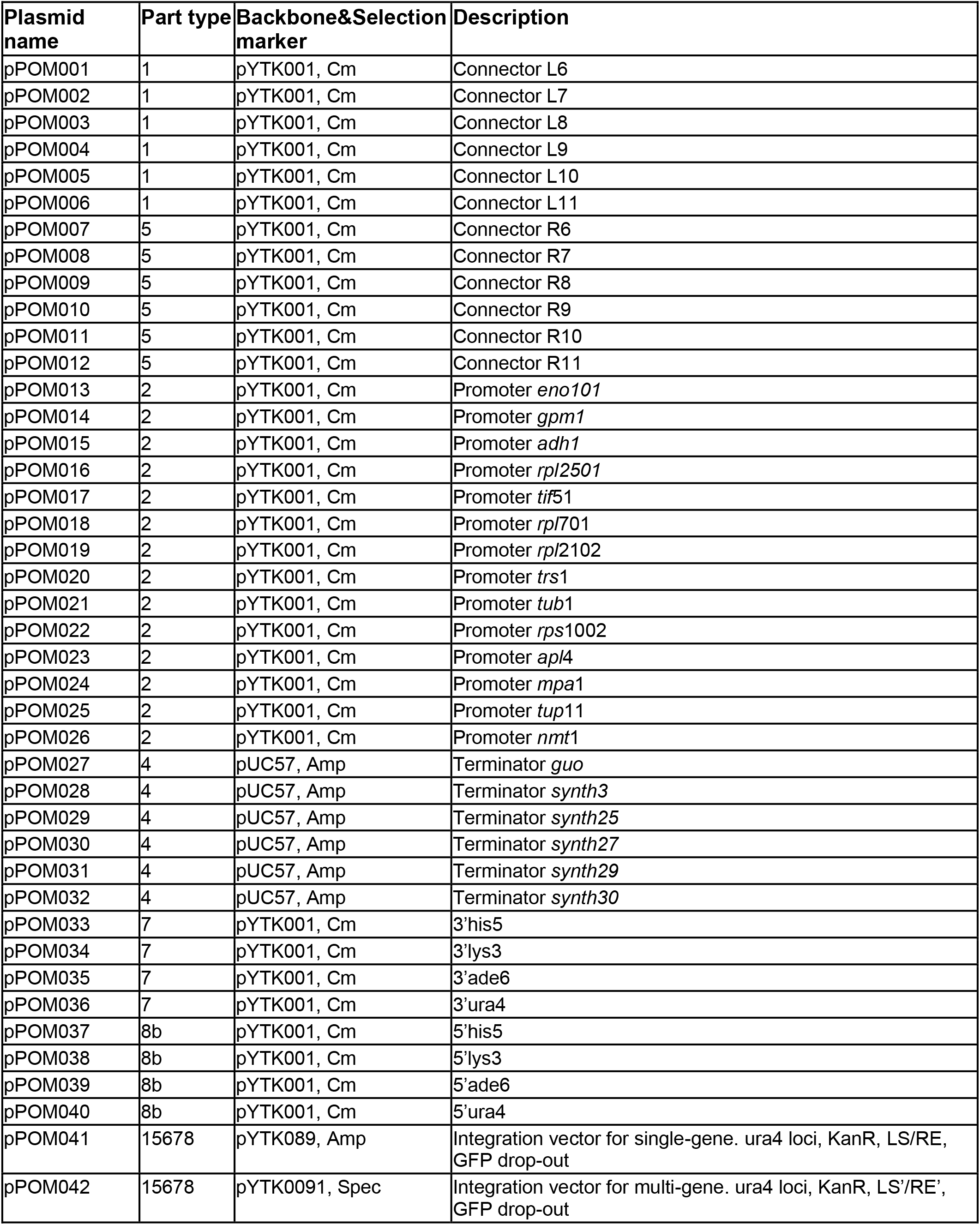
List of part plasmids generated for the toolkit.

**Table S2:**
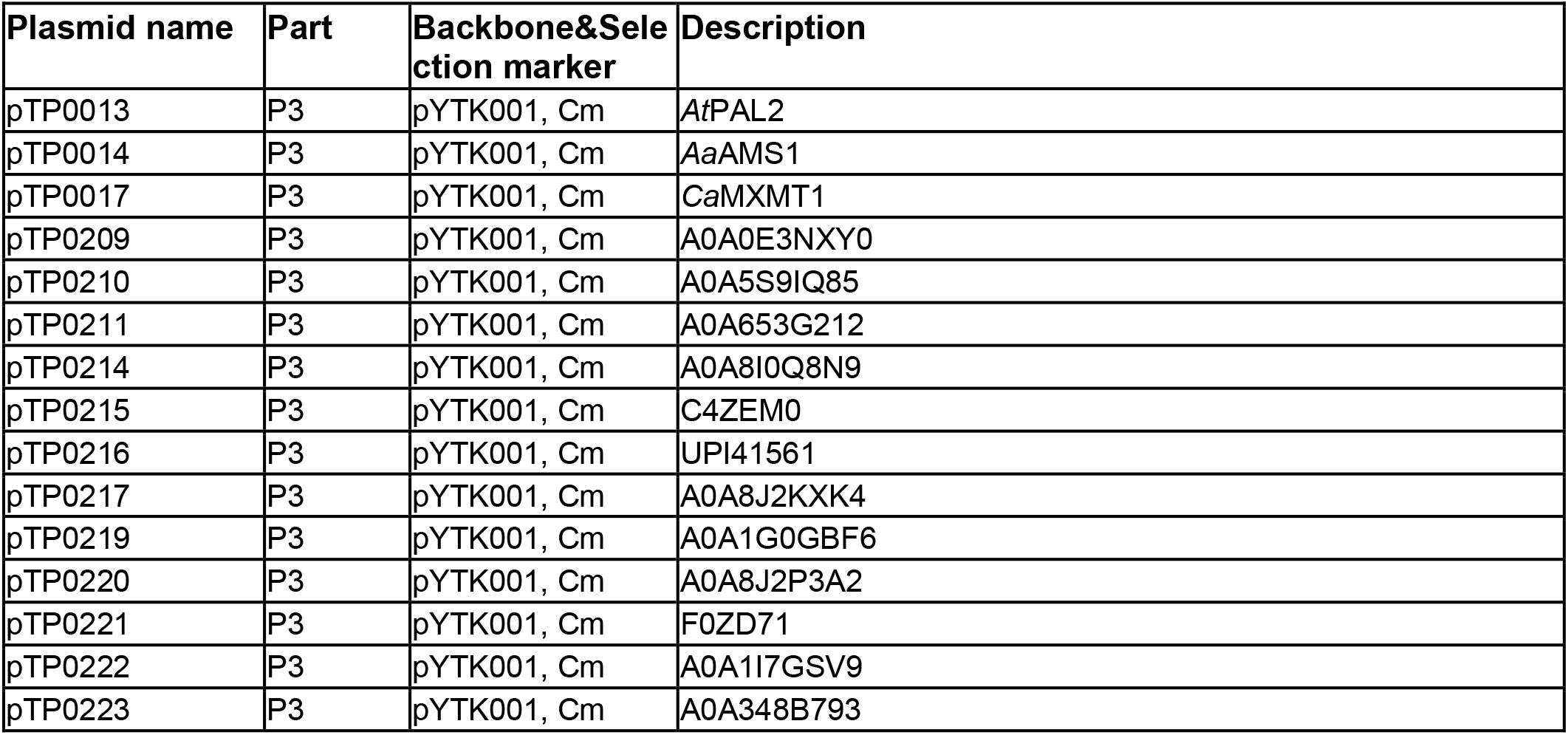
List of part plasmids generated for this study

**Table S3:**
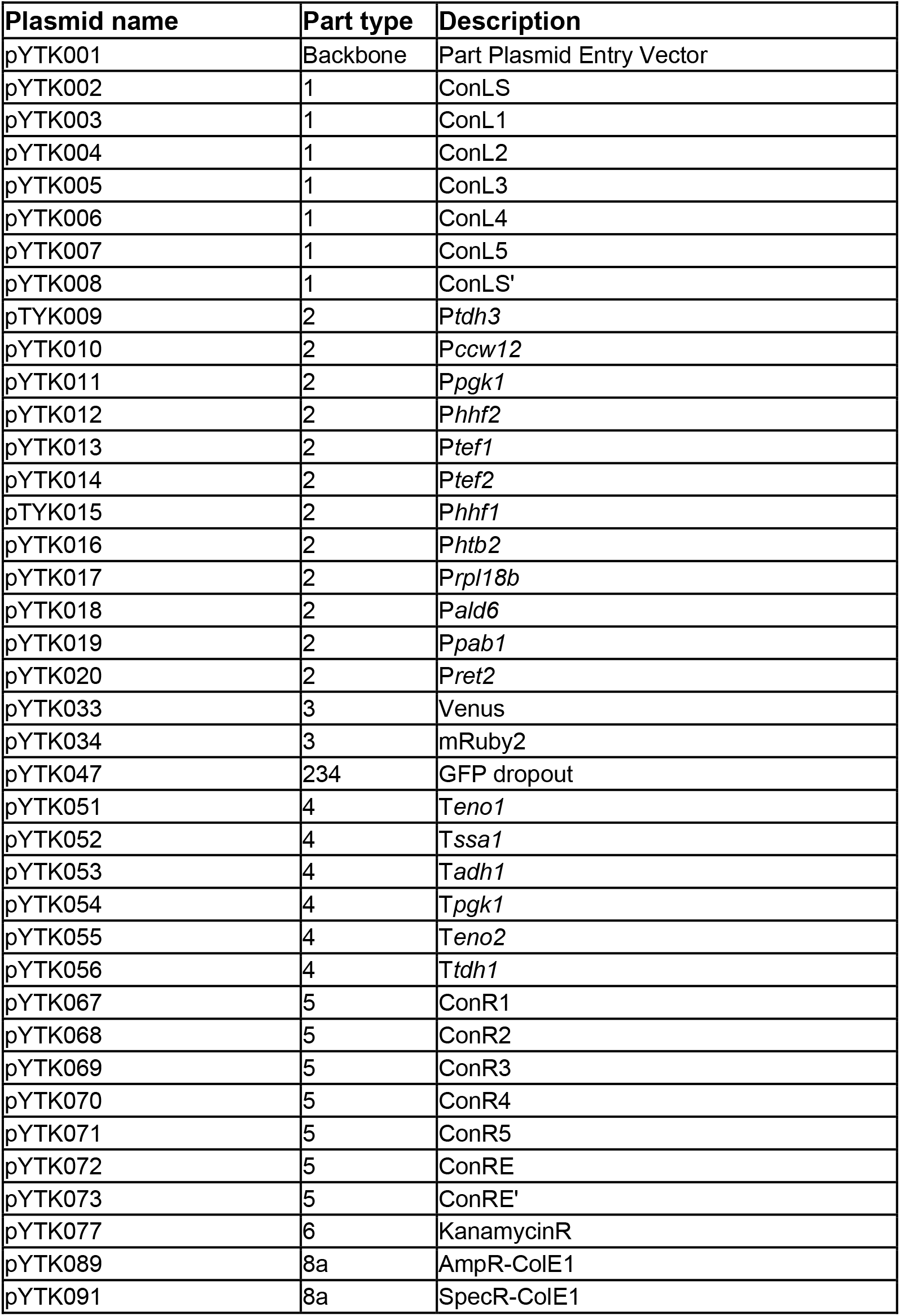
List of part plasmids used in this study.

**Table S4:**
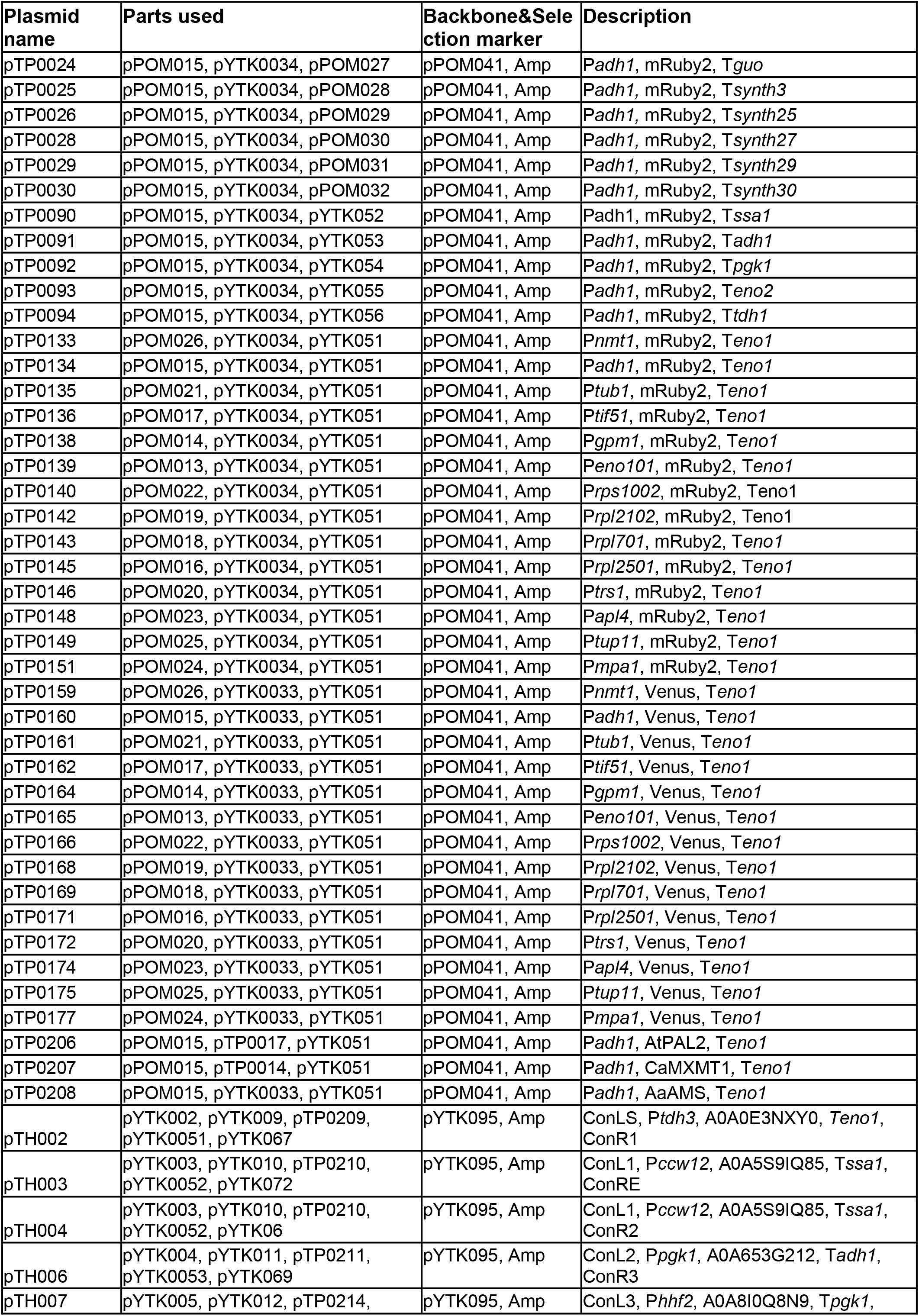
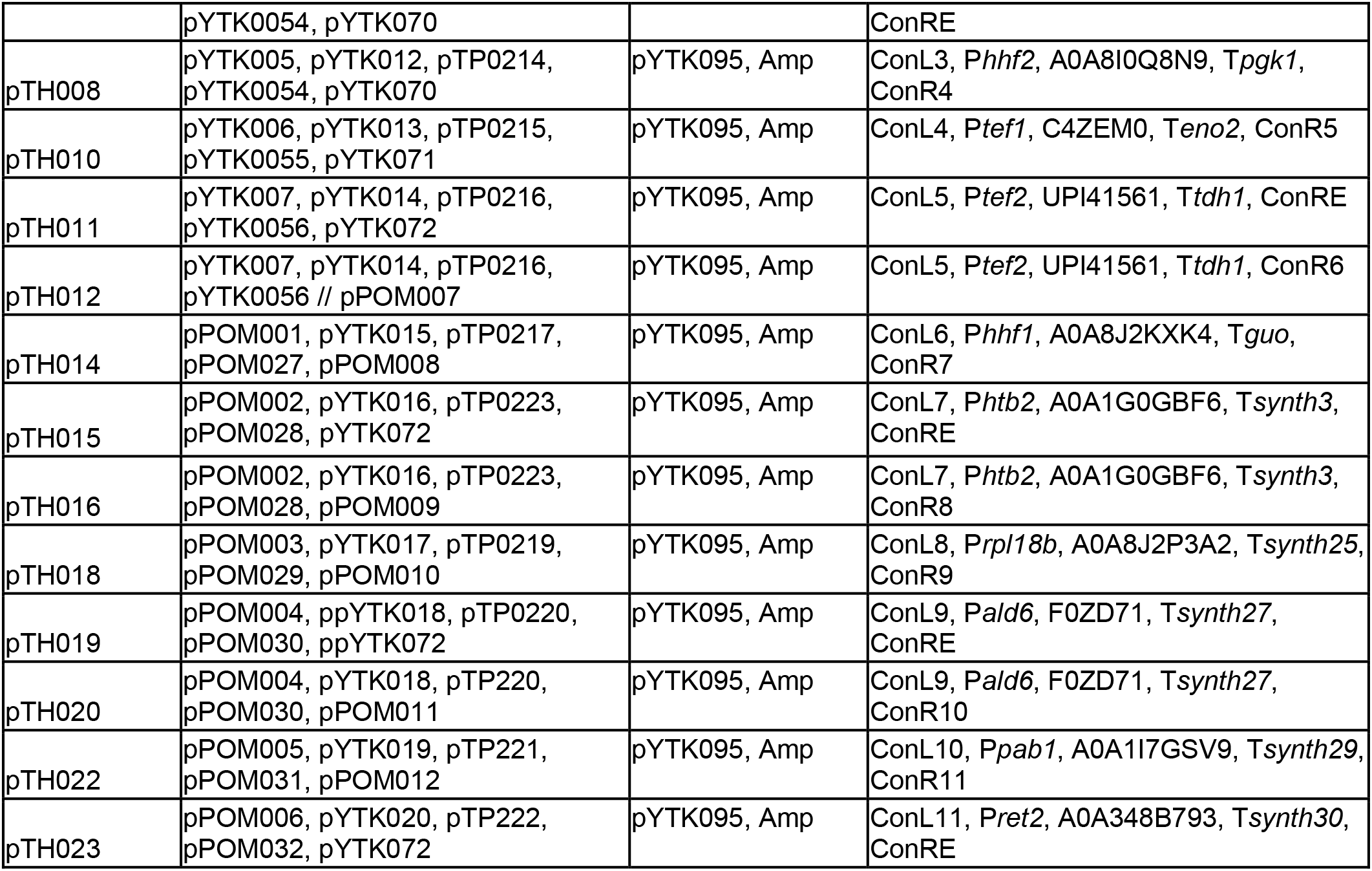
List of single-gene plasmids generated for this study.

**Table S5:**
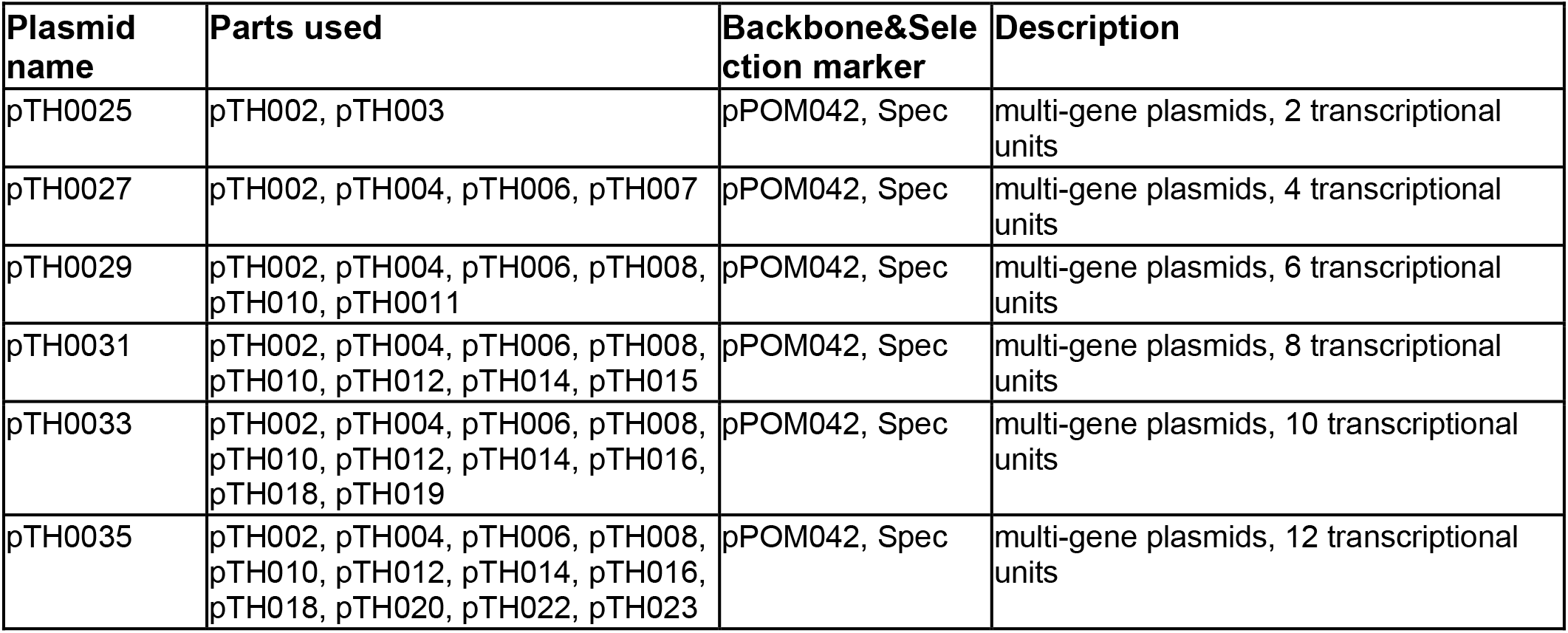
multi-gene plasmids generated for this study

**Table S6:**
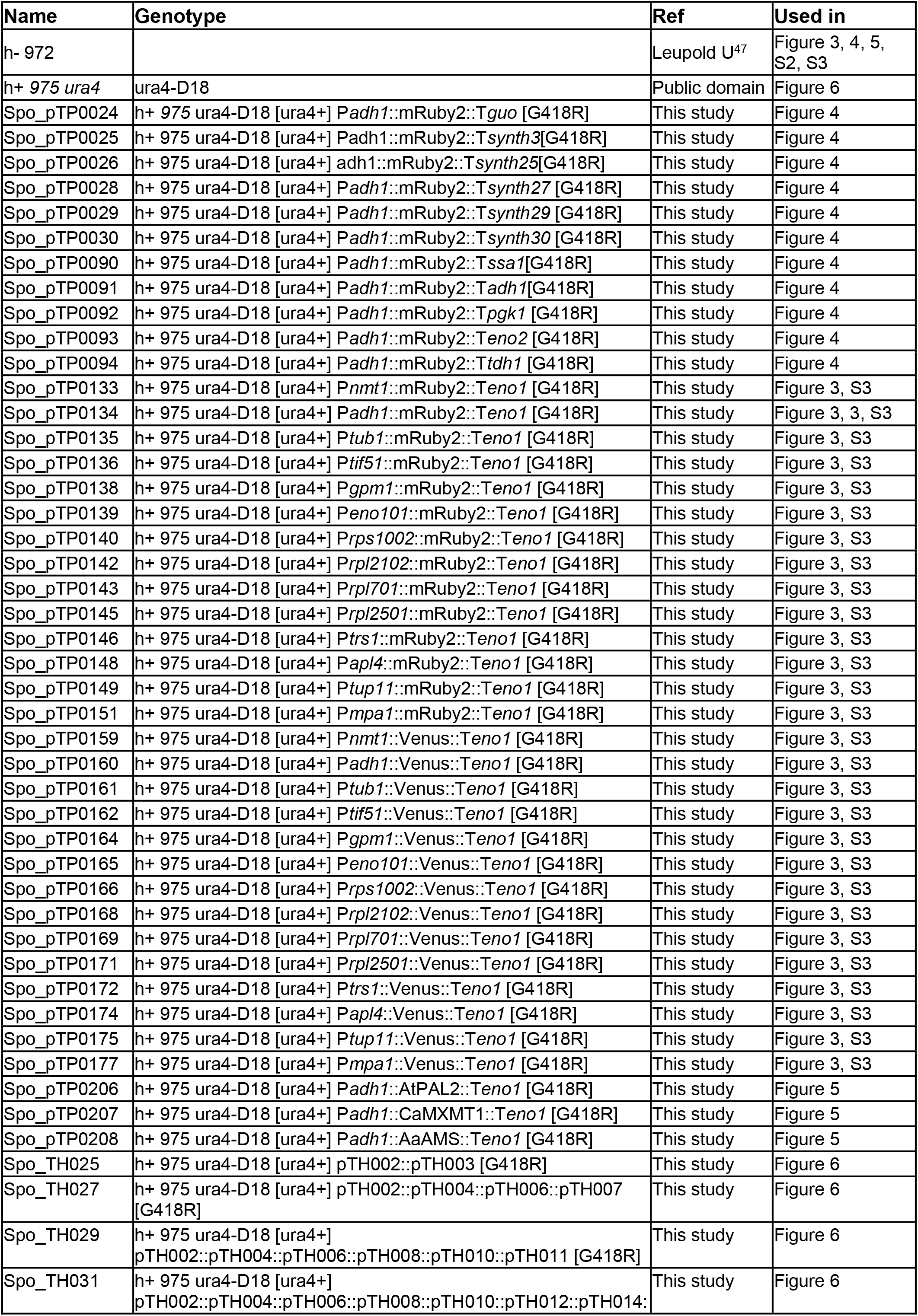
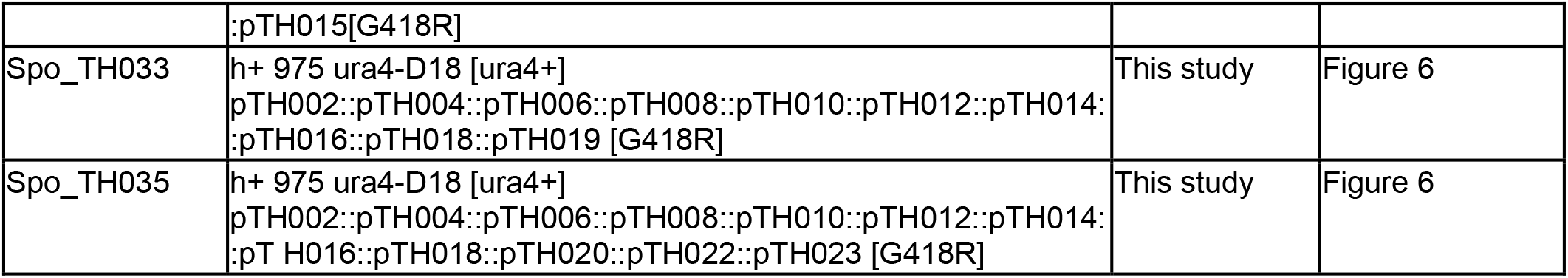
List of strains

**Table S7:**
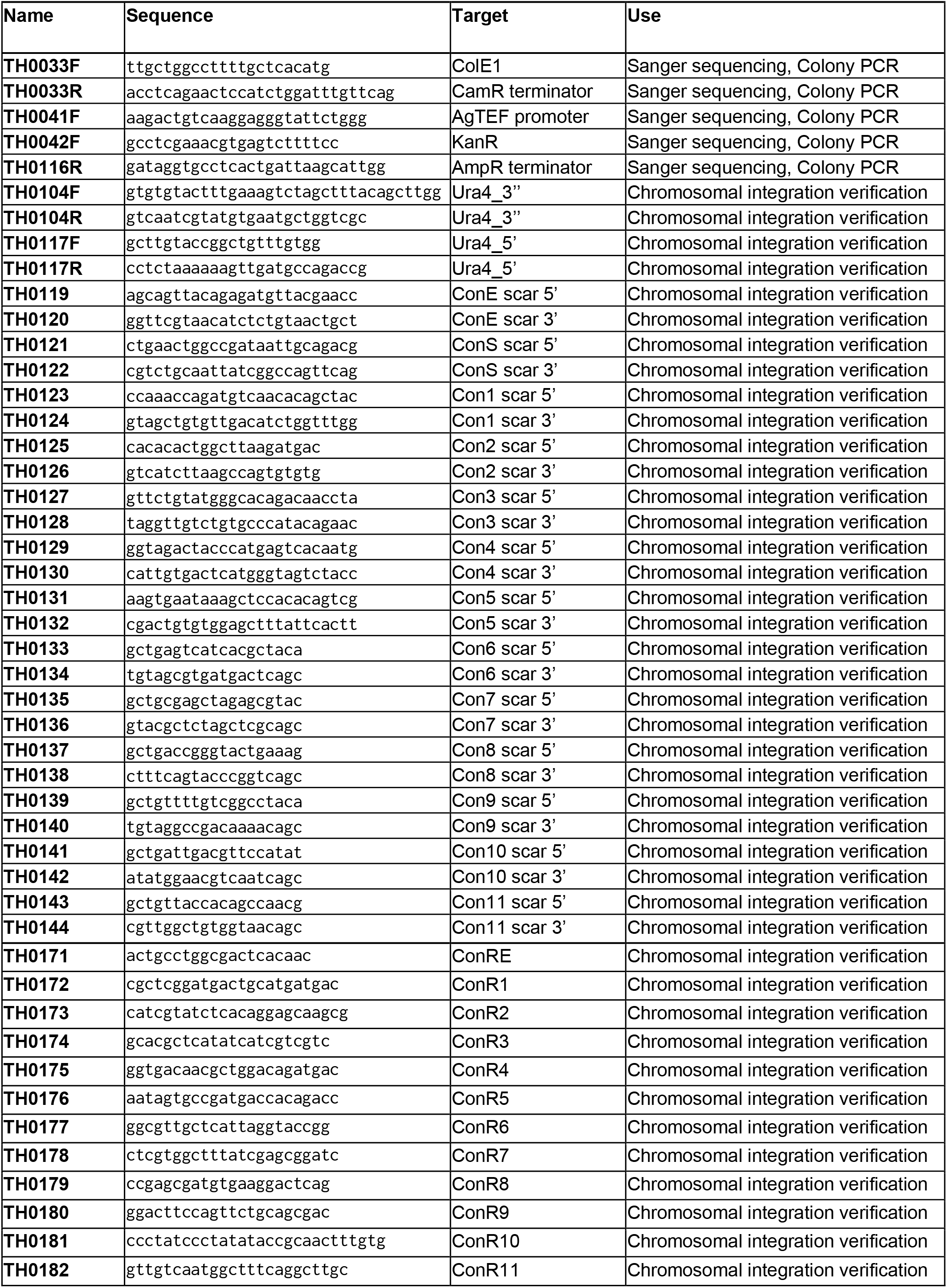
List of genotyping and sequencing primers

**Table S8:**
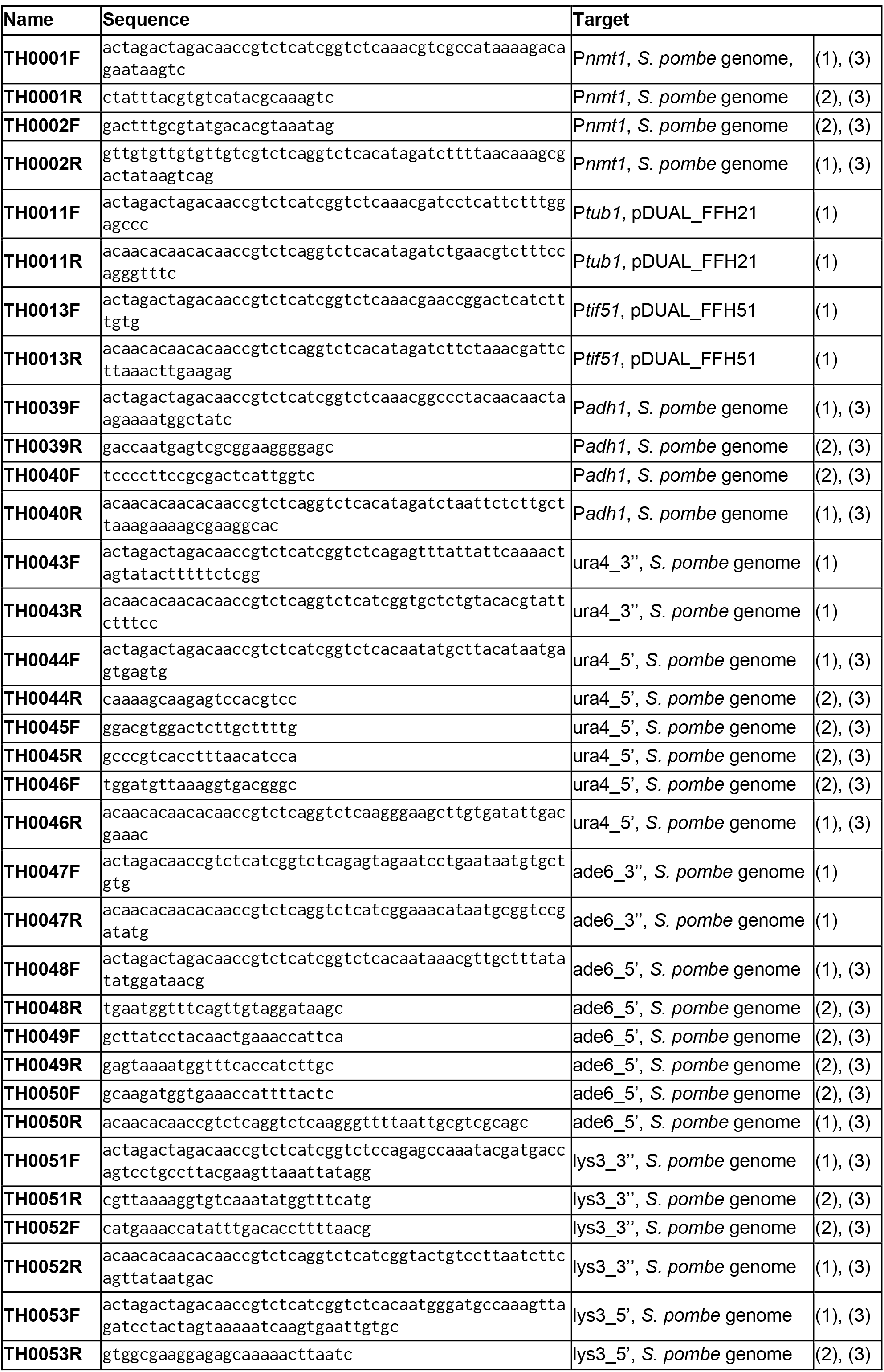
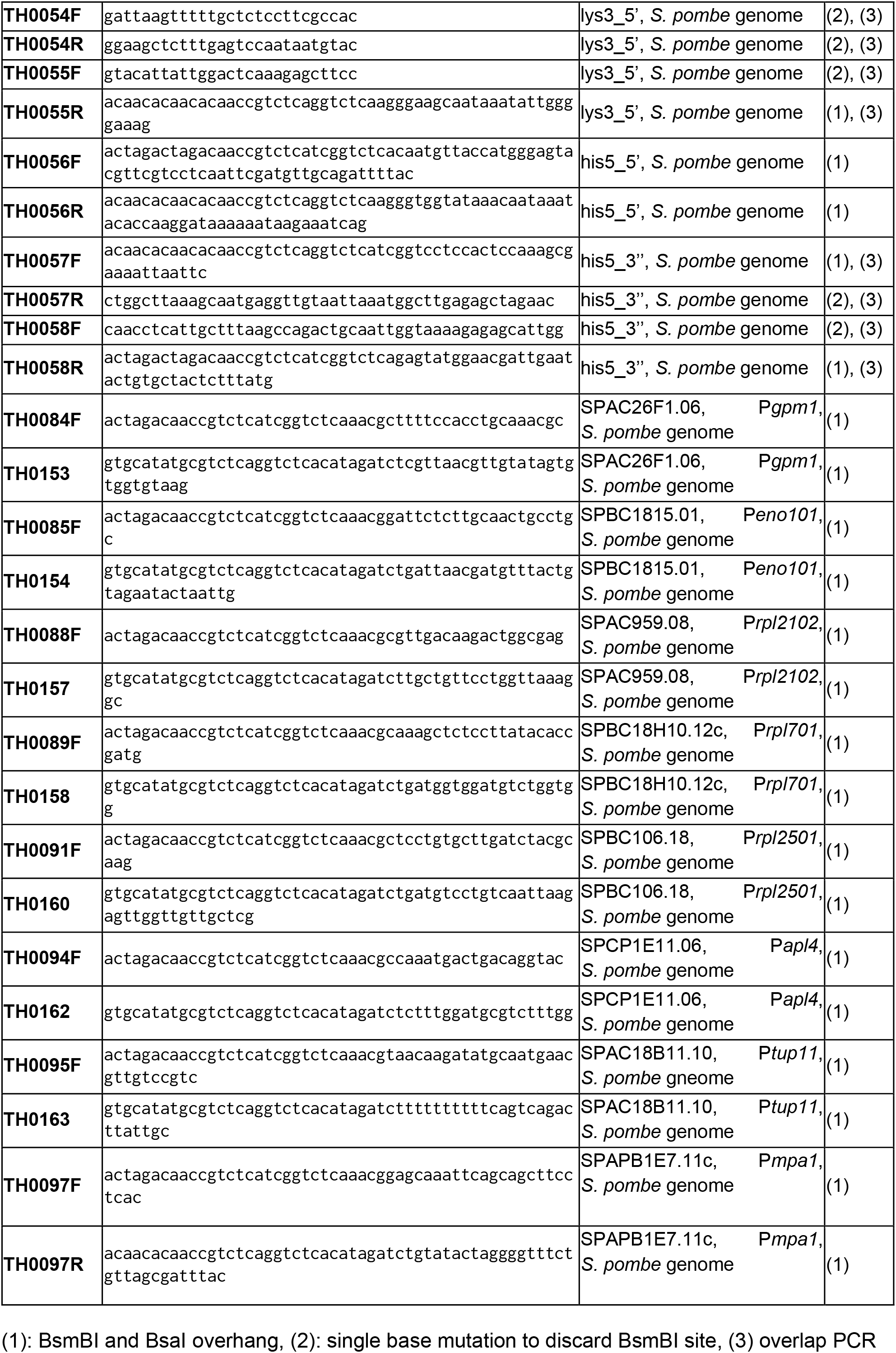
List of primers for DNA part construction

**Table S9:**
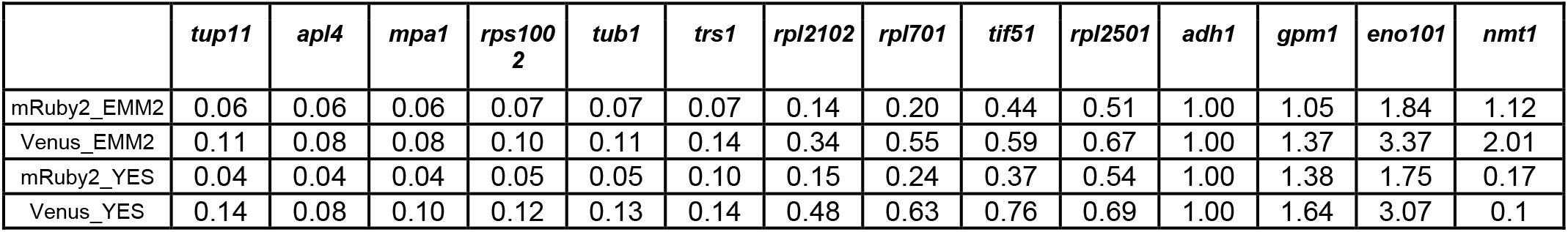
Strength of POMBOX promoters relative to P*adh1*.

